# Lifestyle-associated variation in type IV secretion systems between phytopathogenic and environmental *Ralstonia*

**DOI:** 10.1101/2025.09.26.675681

**Authors:** Tabitha C. Cowell, Matthew L. Cope-Arguello, Gi Yoon Shin, Adam J. Bogdanove, Sara C. D. Carpenter, Kamrun Nahar, Apurba Saha, Tiffany M. Lowe-Power

## Abstract

Type IV secretion systems (T4SSs) are versatile machines with variable functions including DNA uptake and release, protein translocation, and DNA conjugation. However, the diversity, distribution, and functional roles of the T4SS in the *Ralstonia* genus remain poorly understood. The *Ralstonia solanacearum* species complex (RSSC) comprises three species of plant-pathogenic bacteria that cause bacterial wilt disease. The *Ralstonia* genus also includes non-RSSC species that are primarily environmental bacteria and rare opportunistic human pathogens. This study compared the diversity and phylogenetic distribution of T4SSs in the RSSC phytopathogens vs. non-RSSC environmentals. Phylogenetic analysis of VirB4 sequences and synteny analysis revealed 16 distinct T4SS clusters in *Ralstonia*, with ten clusters found in RSSC phytopathogen genomes, twelve in non-RSSC environmental genomes, and six clusters in both groups. Collectively, these gene clusters were more prevalent in non-RSSC environmental genomes. The presence of type IV coupling protein and relaxase genes suggests that at least 14 of these T4SS gene clusters could be putative DNA conjugation systems. The clusters were encoded on accessory plasmids of various sizes or as integrative and conjugative elements (ICEs) on the chromosome or megaplasmid. The putative regions of transfer for T4SS gene clusters in the RSSC phytopathogen genomes often contained type III effectors, type VI secretion toxin/antitoxin clusters, and hemagglutinin gene clusters. In contrast, the non-RSSC environmentals were enriched in heavy metal metabolism and resistance genes. One of the 16 T4SS clusters, cluster i, exhibited evidence of specialization for the RSSC phytopathogens. These findings shed light on the eco-evolutionary differences in the *Ralstonia* genus.

**Impact statement:** The *Ralstonia* genus contains the *Ralstonia solanacearum* species complex (RSSC), a group of globally important plant pathogens that impact food security. These pathogens are known to have an expansive, open pangenome with high levels of gene flow. To shed light on the eco-evolutionary differences between RSSC phytopathogens and closely related environmental species, we explored the potential role of type IV secretion systems (T4SSs) in horizontal gene flow within the *Ralstonia* genus. Surprisingly, these mobile genetic elements were less common in the phytopathogens than the environmentals. Nevertheless, we identified a particular T4SS gene cluster encoded on an accessory plasmid that appears to be specialized to the pathogenic lifestyle of the RSSC. This specialized cluster harbors genes that drive RSSC host range (type III effectors) and others that likely allow these pathogens to antagonize the plant microbiota (type VI secretion toxins and hemagglutinin two partner secretion systems). Moreover, this study sheds light on the cryptic lifestyles of the poorly studied environmental *Ralstonia* species, revealing that their conjugative T4SSs frequently carry heavy metal metabolism and resistance genes. Overall, this study provides a foundation to investigate the functional roles of T4SS clusters and their cargo genes in the fitness of phytopathogenic and environmental *Ralstonia*.

**Data summary:** Supplemental Table S1 lists details of the RSSC phytopathogen, non-RSSC environmental, and Burkholderiaceae family genomes used in this study. Supplemental Table S2A is the final list of 501 VirB4 sequences collectively identified in 636 RSSC phytopathogen genomes and 143 non-RSSC environmental genomes. Supplemental Table S2B includes the annotations and NCBI accessions for every gene and protein sequence in reference clusters a-p. Supplemental Table S2C is the list of putative T4SS cargo genes identified in complete *Ralstonia* genomes. Supplemental Table S2D is the list of putative T4SS cargo genes identified in cluster i regions from draft and complete *Ralstonia* genomes.

Supplemental File S1 is the multiple sequence alignment (MSA) input for HMMER, made from the 753 VirB4 protein sequences from the Burkholderiaceae family genomes. Supplemental Files S2 and S3 are PDF-format versions of the species trees displaying the presence of T4SS gene clusters in RSSC phytopathogens and non-RSSC environmentals, respectively. Supplemental File S4 is the species tree displaying the presence of RSp0179 and RSp1521 in the *Ralstonia* genus. Supplemental File S5 is the MSA input for FastTree 2, made from the VirB4 protein sequences from *Ralstonia* genomes. Supplemental Files S6-S9 are various formats of the VirB4 protein tree in Figure 1. Supplemental Files S10-S25 are the GBK files for reference clusters a-p. Supplemental File S26 is the clinker output comparing the reference clusters. Supplemental Files S27-S36 are the clinker outputs comparing the putative regions of transfer of T4SSs in complete *Ralstonia* genomes. Supplemental File S37 is an MSA analysis comparing RSp1521 to AcvB homologs. Supplemental File S38 is the clustered average nucleotide identity (ANI) matrix for the putative regions of transfer of cluster i T4SSs in draft and complete *Ralstonia* genomes. Supplemental File S39 is the clinker output comparing the cluster i regions in draft and complete *Ralstonia* genomes. All Supplemental Tables and Files are available through Zenodo (DOI: 10.5281/zenodo.14873518).

This study’s newly sequenced genomes, Gazipur 4 and Gazipur 5, are available on NCBI with the following accessions: GCF_049860735.1 and GCF_049860725.1, respectively. The two contigs containing the cluster j T4SS were submitted to NCBI through the Third Party Annotation (TPA) section of the DDBJ/ENA/GenBank databases with the following accessions: BK072068-BK072069. These contigs were assembled from SRA reads under the accession: SRR18649448.

## Introduction

Each environment presents specific opportunities and challenges for bacteria. For example, soil-dwelling bacteria must survive unpredictable nutrient and terminal electron acceptor availability, variations in moisture, and intermicrobial competition. Host-colonizing bacteria must survive many of those challenges as well as host immune factors, such as bursts of reactive oxygen species and antimicrobial molecules. To seize the opportunities and overcome the challenges of their lifestyles, bacteria adapt through evolution where selection acts on pre-existing genetic variation derived from *de novo* mutations or horizontal gene transfer (HGT). Cases have been identified in which certain strains of an environmental bacterial species become pathogenic due to the presence of horizontally acquired DNA (1–5). Horizontally transferred DNA can include conjugative plasmids, integrative and conjugative elements (ICEs), non-conjugative plasmids, transposons, and phages (6,7).

By definition, conjugative plasmids and ICEs are self-transmissible because they encode conjugative molecular machines called type IV secretion systems (T4SSs) (8–10). Additionally, these mobile genetic elements often carry non-T4SS genes that confer new functions that may enable niche adaptation. A common marker of T4SSs is the VirB4 protein sequence because VirB4 family proteins are highly conserved ATPases that are associated with all known T4SSs (10–13). This family includes VirB4, TrbE, CagE, TraC, TraU, IcmB, DotO, and Tfc16. Determining that a given T4SS gene cluster encodes a conjugation machine requires experimental validation because in addition to DNA conjugation, T4SSs may carry out DNA uptake, environmental release of DNA, or protein translocation (8,10,14). Nevertheless, the presence of genes encoding the type IV coupling protein (T4CP) and relaxase can provide circumstantial evidence that the T4SS is conjugative. The T4CP recruits the DNA and/or protein substrate(s) to the membrane-bound T4SS complex (8,15). A T4CP is required for DNA conjugation (15,16) and DNA release systems (10), and is usually, but not always, associated with protein translocation systems (10,13). Additionally, a relaxase is required for DNA conjugation (10,13,15,17) and DNA release systems (10,13), although the relaxase is not considered a true T4SS component. Therefore, the presence of a T4CP gene in a T4SS gene cluster and a relaxase gene in the vicinity is an indicator that the T4SS in question may function as a DNA conjugation or DNA release system. However, the absence of an identifiable T4CP or relaxase gene cannot be used to rule out these potential functions.

A model genus to explore the question of niche adaptation of pathogenic versus environmental lifestyles is *Ralstonia*. The *Ralstonia* genus is composed of two major clades: (i) the aggressive plant pathogens in the *Ralstonia solanacearum* species complex (RSSC) and (ii) the environmental bacteria isolated from diverse habitats. Hereafter, we call these groups the RSSC phytopathogens and the non-RSSC environmentals, respectively. The RSSC phytopathogens are the primary causative agent of bacterial wilt disease and include three species: *Ralstonia solanacearum*, *R. pseudosolanacearum*, and *R. syzygii* (18). Most RSSC phytopathogens are environmentally transmitted through soil and surface water between infection cycles where they colonize the rhizosphere, xylem, and apoplastic spaces (19). The non-RSSC environmentals include at least seventeen species: *R. chuxiongensis*, *R. condita*, *R. edaphi*, *R. flaminis*, *R. holmesii*, *R. insidiosa*, *R. mannitolilytica*, *R. mojiangensis*, *R. pickettii*, *R. psammae*, *R. thomasii*, *R. wenshanensis*, and five unnamed *Ralstonia* spp. Less is known about the ecological niches of non-RSSC environmentals, but they have been isolated from more diverse natural and human-made habitats: plant rhizosphere, soil, surface water, built water infrastructure, filter-sterilized pharmaceuticals, and opportunistic infections of physiologically and immunocompromised patients. Although the environmentals can be opportunistic pathogens, these are rare outcomes, largely resulting from colonization of indoor plumbing and contamination of pharmaceuticals (20).

T4SSs are widespread and can be found in many bacterial and archaeal taxa (8,10,12,14), and are often studied for their contribution to behavior and evolution of pathogens (1,4,21). Few studies have compared T4SS biology between well-established pathogens and their environmental relatives. Here, we elucidate the diversity and phylogenetic distribution of T4SSs within RSSC phytopathogens and non-RSSC environmentals. We used VirB4 protein sequences to identify T4SS gene clusters in the *Ralstonia* pangenome. We characterized the gene clusters by synteny and sequence diversity, compared their phylogenetic distributions within the genus, and inferred their adaptive potential.

## Methods

### Genomes and accessions

Supplemental Table S1 lists details of the RSSC phytopathogen, non-RSSC environmental, and Burkholderiaceae family genomes used in this study. The quality of all *Ralstonia* genomes was assessed using CheckM (22) in KBase (23) and only genomes with greater than 96% completeness and less than 2.5% contamination were included. Bioinformatic analysis used sets of genomes: 394 RSSC phytopathogen genomes available as NCBI assemblies in 2023, 636 RSSC phytopathogen genomes available as NCBI assemblies or as raw reads on NCBI SRA in 2024, 143 non-RSSC environmental genomes from 17 genomospecies available as NCBI assemblies in 2023, and 923 Burkholderiaceae family genomes that include the 394 RSSC phytopathogen genomes and 529 genomes from at least 252 genomospecies. Most genomes were downloaded from the NCBI assembly database, but 145 RSSC phytopathogen genomes, mostly from Greenrod *et al.* (2022) (24), were assembled from raw reads acquired from NCBI SRA. Genome assembly was performed on KBase; reads were trimmed with Trimmomatic (Illumina) (25) or Filtlong (Nanopore) (26), assembled with SPAdes (Illumina) (27) or Unicycler (hybrid Nanopore and Illumina) (28), and annotated with Prokka (29). These assemblies can be accessed from a public KBase narrative (https://narrative.kbase.us/narrative/189849). The two contigs containing the cluster j T4SS were submitted to NCBI through the Third Party Annotation (TPA) section of the DDBJ/ENA/GenBank databases with the following accessions: BK072068-BK072069. These contigs were assembled from SRA reads under the accession: SRR18649448.

We isolated two *Ralstonia pseudosolanacearum* strains from *Solanum melongena* stems in Gazipur, Bangladesh. Each strain was grown on SMSA agar plates (1 g/L casamino acids, 10 g/L peptone, 5 mL/L glycerol, 15 g/L agar, 50 mg/L 2,3,5-triphenyltetrazolium chloride, 5 mg/L crystal violet, 100 mg/L polymyxin B sulphate, 25 mg/L bacitracin A, 5 mg/L chloramphenicol, and 0.5 mg/L penicillin G) (30) and single colonies were inoculated into 6 mL of CPG broth (1 g/L casamino acids, 10 g/L peptone, 5 g/L glucose, and 1 g/L yeast extract) (31). Liquid cultures were grown overnight in a shaking incubator at 250 rpm and 28°C. We extracted gDNA using 2 mL of culture and the MasterPure Gram Positive DNA Purification Kit (Biosearch Technologies, Cat. No. MGP04100) using the manufacturer’s protocol with doubled reagent volumes. The genomes of these strains were sequenced by Plasmidsaurus using Oxford Nanopore Technology and assembled with Flye v2.9.1 (32). The genome sequences for Gazipur 4 and Gazipur 5 are available on NCBI with the following accessions: GCF_049860735.1 and GCF_049860725.1, respectively.

### Synteny analysis

We used clinker (33) to visualize synteny of gene neighborhoods with a threshold of 30% global amino acid identity.

### Identification of VirB4 protein sequences using a custom hidden Markov model

Initial BLASTp (34) searches for VirB4 sequences in RSSC phytopathogen genomes revealed there were at least six VirB4 variants, so we used a hidden Markov model (HMM) approach in the KBase platform to robustly identify divergent VirB4 protein sequences. Briefly, we BLASTp searched the six VirB4 proteins against 923 Burkholderiaceae family genomes, used the hits to build a preliminary HMM, searched the preliminary HMM against the 923 Burkholderiaceae family genomes, built a final HMM from the hits, and searched the final HMM against 636 RSSC phytopathogen and 143 non-RSSC environmental genomes. We queried six divergent VirB4 protein sequences against the 923 Burkholderiaceae family genomes using BLASTp with 20% identity and 20% alignment overlap thresholds (WP_011002496.1, WP_003274595.1, WP_042568051.1, WP_247391909.1, WP_230646236.1, and WP_230645232.1). The 617 combined hits were aligned in a multiple sequence alignment (MSA) using MUSCLE (35) with a maximum of 16 iterations. With HMMER (36), this MSA was queried against the 923 Burkholderiaceae family genomes with thresholds of 20% model coverage and a bit score of 20, yielding 753 VirB4 sequences. Using the previously described parameters, the 753 VirB4 protein sequences were aligned using MUSCLE and the resulting MSA (Supplemental File S1) was queried against 636 RSSC phytopathogen and 143 non-RSSC environmental genomes using HMMER. The final list of 496 VirB4 sequences identified in those genomes is included in Supplemental Table S2A. We used clinker to analyze the gene neighborhoods of a subset of the identified *virB4* genes to verify that they were near other T4SS genes.

Genes annotated as pseudogenes are missed by BLASTp and HMMER searches. So, in addition to the 496 VirB4 sequences identified by the HMMER search, we manually identified five T4SS gene clusters with *virB4* genes that were annotated as pseudogenes. These are included in the list in Supplemental Table S2A.

### Construction of species trees and protein trees

Species trees and protein trees were constructed on the KBase platform (23). Newick files were visualized in iTOL (37). All species trees (Supplemental Files S2-S4) were built using the KBase “Insert Set of Genomes Into SpeciesTree” pipeline, which uses protein sequences from 49 conserved genes (COG0012 (YchF), COG0013 (AlaS), COG0016 (PheS), COG0018 (ArgS), COG0030 (KsgA), COG0041 (PurE), COG0046 (PurL), COG0048 (RpsL), COG0049 (RpsG), COG0051 (RpsJ), COG0052 (RpsB), COG0072 (PheT), COG0080 (RplK), COG0081 (RplA), COG0082 (AroC), COG0086 (RpoC), COG0087 (RplC), COG0088 (RplD), COG0089 (RplW), COG0090 (RplB), COG0091 (RplV), COG0092 (RpsC), COG0093 (RplN), COG0094 (RplE), COG0096 (RpsH), COG0097 (RplF), COG0098 (RpsE), COG0099 (RpsM), COG0100 (RpsK), COG0102 (RplM), COG0103 (RpsI), COG0105 (Ndk), COG0126 (Pgk), COG0127 (RdgB), COG0130 (TruB), COG0150 (PurM), COG0151 (PurD), COG0164 (RnhB), COG0172 (SerS), COG0185 (RpsS), COG0186 (RpsQ), COG0215 (CysS), COG0244 (RplJ), COG0256 (RplR), COG0343 (Tgt), COG0504 (PyrG), COG0519 (GuaA), COG0532 (InfB), COG0533 (YgjD)). The KBase SpeciesTree pipeline uses a curated MSA for each COG (Clusters of Orthologous Groups) where poorly aligned sections of the MSAs are trimmed using GBLOCKS (38), the trimmed MSAs are concatenated, and an approximately-maximum-likelihood tree is constructed using FastTree 2 (39) with the -fastest setting. Protein trees were built by constructing MSAs with MUSCLE with a maximum of 16 iterations before building approximately-maximum-likelihood protein trees with FastTree 2 using default settings in KBase (Supplemental Files S5-S9).

### Gene annotation of T4SS gene clusters

We selected one member of each of the 16 *Ralstonia* T4SS gene clusters to be reference clusters. For clusters with only one member, that member was selected as the reference cluster. For clusters with multiple members, the reference cluster was selected from a genome using the following priorities: availability on NCBI RefSeq, a chromosome-scale complete assembly, containing more than one T4SS gene cluster, and being from a model lab strain.

To annotate T4SS genes in the T4SS reference gene clusters, we used three annotation sources: NCBI Prokaryotic Genome Annotation Pipeline (PGAP) (40), NCBI Conserved Domain Database (CDD) (41), and MacSyFinder v2 with CONJscan models (42,43). We ran MacSyFinder using the “unordered” parameter, which ignores gene order to account for draft genome assemblies, with a FASTA file containing all amino acid sequences in a given genome. For the CONJscan models, we used both the “Chromosome all” and “Plasmids all” parameters in separate runs. When annotations conflicted, priority was given to MacSyFinder/CONJscan, then NCBI RefSeq/GenBank, then NCBI Conserved Domain Database. Operon-mapper (44) was used to predict operons in genomes used to build the reference T4SS gene clusters. Any predicted operons that contained an annotated T4SS gene were included in the reference cluster. NCBI accessions for genomes, genes, and proteins for every gene in reference clusters a-p can be found in Supplemental Table S2B and Supplemental Files S10-S26.

To broadly classify the T4SS gene clusters, we used the nomenclature established by the F/P/I/GI-type classification system (10,11). However, the CONJscan models are based on the different mating pair formation (MPF) classification system and include MPF_F_, MPF_T_, MPF_I_, MPF_G_, MPF_B_, MPF_C_, MPF_FA_, and MPF_FATA_ models (13,42,45). Between these two classification systems, MPF_F_ corresponds to F-type, MPF_T_ to P-type, MPF_I_ to I-type, and MPF_G_ to GI-type. The remaining MPF classes include T4SSs that are not represented by the F/P/I/GI-type classes.

### Determining the phylogenetic distribution of T4SS gene clusters using the VirB4 protein sequences

To determine the phylogenetic distribution of the T4SS gene clusters in *Ralstonia* (clusters a-p as described in the Results), we used BLASTp with 40% sequence identity and 50% alignment overlap thresholds to search 636 RSSC phytopathogen genomes and 143 non-RSSC environmental genomes for each of the 16 identified VirB4 protein sequences, one from each of the reference gene clusters. The NCBI accessions for the VirB4 protein or gene sequences of each reference cluster and the genome it came from are as follows: WP_011002496.1, GMI1000 (a); WP_247391909.1, T25_UW811 (b); WP_214372649.1, SY1 (c); WP_123203755.1, Tg03 (d); WP_230645232.1, LMG10661 (e); WP_003274595.1, MolK2 (f); WP_165858801.1, SL1931 (g); WP_230646236.1, LMG10661 (h); WP_042568051.1, UW163 (i); YO100T327_00165, YO100_T327 (j); WP_103518556.1, ICMP-8657 (k); WP_009242089.1, WCHRI065437 (l); WP_182594753.1, s11 (m); WP_004630343.1, WCHRI065162 (n); WP_354534808.1, 1138 (o); and WP_024979717.1, SSH4 (p). There was one exception for cluster d, which has two members with 24.97% identity and 92% coverage by BLASTp comparison of their VirB4 protein sequences. Therefore, the 40% sequence identity threshold was too stringent for this particular case. We used the iTOL annotation editor for Google Sheets (37) and the results from BLASTp to annotate the RSSC phytopathogen and non-RSSC environmental species trees (Supplemental Files S2-S3). Please note that the genomes SY1 and LMG10661 are suppressed by NCBI. However, the protein sequence accessions provided are still accessible through NCBI.

### Identifying putative region of transfer and putative cargo genes

The analysis of putative cargo genes was only performed for T4SS gene clusters from complete genomes, so clusters b, h, j, k, o, and p were not analyzed due to a lack of complete genomes. For the RSSC phytopathogens, 33 putative regions of transfer from 25 complete genomes were analyzed. For the non-RSSC environmentals, 33 putative regions of transfer from 14 complete genomes were analyzed. For T4SS gene clusters encoded on accessory plasmids, the putative region of transfer was considered to be the entire plasmid, except for NZ_CP146093.1, which was an accessory plasmid with two T4SS clusters. For NZ_CP146093.1 and for T4SS gene clusters encoded on the chromosome or megaplasmid, the putative region of transfer was determined by manual inspection using synteny analysis with clinker. During manual inspection, regions were determined by identifying genes that consistently acted as a boundary, such as an integrase gene for cluster a, and/or by identifying a consistent insertion location characteristic, such as a tRNA gene for cluster c. See Supplemental Files S27-S36 for clinker output files with putative regions of transfer from each cluster.

Within each putative region of transfer, we searched for type III secretion system (T3SS) effector genes from the Ralsto T3E database (46) using BLAST+ with a threshold of 40% amino acid identity (34) followed by manual verification of each hit. We also searched for type VI secretion system (T6SS) *vgrG*-linked auxiliary clusters. We have previously curated 25 *vgrG*-linked auxiliary clusters from RSSC phytopathogen genomes (47) and we curated 19 additional *vgrG*-linked auxiliary clusters from non-RSSC environmental genomes using the method described in Aoun and Georgoulis *et al*. (2024). We first used clinker to search the putative regions of transfer for *vgrG*-linked auxiliary clusters based on a global amino acid identity threshold of 30%. As a second approach, we performed BLASTp searches with seven representative VgrG alleles with 20% local identity and 20% coverage thresholds. The representatives were chosen using clinker to identify VgrG alleles with less than 30% global amino acid identity between each other. We searched for hemagglutinin two-partner secretion (TPS) clusters using BLASTp followed by manual verification of each hit, regardless of percent identity and percent coverage (query sequences: WP_240442336.1, WP_118891273.1, and WP_118891272.1). Additionally, we searched for metabolism-related genes using DRAM with a bit score threshold of 60 (48), dbCAN2 with an e-value threshold of 1e-15 (49), and ModelSEED v2 with sequential gapfilling (50) in KBase and METGeneDb v2 (51) using BLAST+ with a 40% identity threshold. We combined the results of the four different metabolism gene searches, then divided them into two categories: (i) heavy metal metabolism and resistance genes and (ii) other metabolism genes. Finally, we used MOBscan (52) to search for relaxase genes. MOBscan uses a protein sequence-based HMM profile for each of the following MOB families: MOB_P1_, MOB_P2_, MOB_P3_, MOB_Q_, MOB_H_, MOB_C_, MOB_V_, MOB_B_, MOB_F_, MOB_T_, and MOB_M_.

### Average nucleotide identity of putative regions of transfer

Average nucleotide identity (ANI) of putative regions of transfer was calculated using pyANI-plus v0.0.1 (53) with the ANIb method, which uses BLAST+. Hierarchical clustering of the ANI matrix was performed using Morpheus (54) with the Euclidean distance metric.

### Statistical analysis

All graphing and statistical analyses were performed in GraphPad Prism v10.4.1 for Windows.

## Results

### There are sixteen distinct T4SS gene clusters in the *Ralstonia* genus

To broadly identify T4SSs in *Ralstonia*, we used VirB4 as a marker. We built a hidden Markov model (HMM) from 753 VirB4 protein sequences identified by preliminary BLASTp searches against 923 Burkholderiaceae genomes. Querying the HMM against 636 RSSC phytopathogen and 143 non-RSSC environmental genomes identified 496 VirB4 sequences. A protein tree of the 496 sequences revealed at least 13 clades. To test the hypothesis that there were multiple, distinct T4SS gene clusters, we analyzed the synteny and global amino acid sequence identity of the *virB4* gene neighborhoods using clinker (33). In most cases, gene organization patterns reinforced the phylogenetic groupings, but in two cases, synteny conservation within a cluster and differences in synteny between the clusters allowed us to differentiate and divide the clusters (b vs. k and h vs. p). Through this combined phylogenetic and syntenic approach, we ultimately identified 16 distinct T4SS gene clusters in the *Ralstonia* genus, which we referred to as clusters a-p (**Figure 1**).

**Figure 1.**
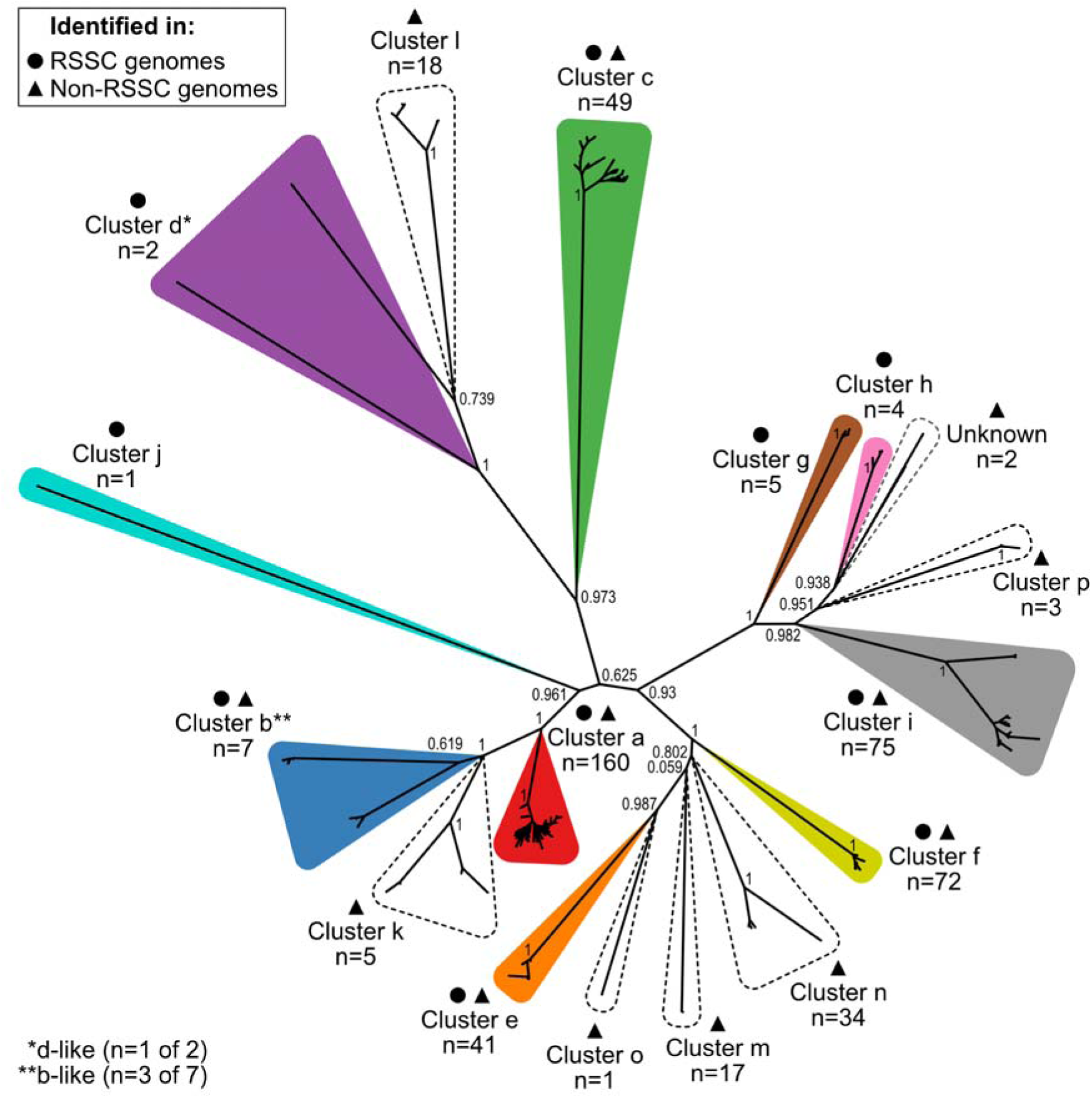
Sequence diversity of the T4SS marker VirB4 in the *Ralstonia* genus. VirB4 protein sequences (n=496) were identified by a custom hidden Markov model (HMM) search, and an approximately-maximum-likelihood tree was built with FastTree 2 from a MUSCLE multiple sequence alignment (MSA) of the VirB4 sequences. Bootstrap values (0 to 1) from 1,000 resamples are shown for key nodes. The unrooted tree was visualized in iTOL and labeled using Affinity Designer. The 16 distinct T4SS gene clusters (a-p) are outlined and labeled. Circles denote clusters that are present in the RSSC phytopathogens and triangles denote clusters that are present in the non-RSSC environmentals. Colored clusters were found in RSSC phytopathogen genomes, and clusters with dashed outlines were not found in the RSSC phytopathogens. Supplemental files contain the *virB4* gene list (Supplemental Table S2A), the VirB4 protein sequence MSA (Supplemental File S5), the Newick file for the tree (File S6), a PDF of the unrooted tree with branch labels (File S7), and PDFs of the rectangular tree with branch labels, without and with colors (Files S8 and S9).

Two clusters, b and d, contain members that are better described as “b-like” and “d-like”, respectively. This is due to the members branching closely to a named cluster in the VirB4 tree (**Figure 1**) and sharing conserved gene organization, but having less than 30% global pairwise amino acid identity for several genes in the cluster (Supplemental Figure S1). For downstream analyses, the three b-like clusters were considered members of cluster b, and the one d-like cluster was considered a member of cluster d. Additionally, we identified two *virB4* genes in two non-RSSC phytopathogen genomes that were not able to be syntenically matched to a particular gene cluster due to an inadequate number of T4SS genes nearby and less than 30% global amino acid identity for most genes present (Supplemental Figure S1). These VirB4 sequences branched close to clusters h and p, and we describe them as “Unknown” (**Figure 1**). For downstream analyses, these *virB4* genes were used whenever we did not differentiate by cluster type.

The clusters varied in their prevalence in the *Ralstonia* genus as well as in their prevalence in the RSSC phytopathogens vs. the non-RSSC environmentals. The most commonly identified T4SS clusters were cluster a (n=160), cluster i (n=75), and cluster f (n=72), while the least commonly identified were cluster j (n=1), cluster o (n=1), and cluster d (n=2) (**Figure 1**). Six T4SS gene clusters were found in both RSSC phytopathogen and non-RSSC environmental genomes (clusters a, b, c, e, f, and i). Four T4SS clusters were only observed in RSSC phytopathogen genomes (clusters d, g, h, and j) while six T4SS clusters were only observed in the non-RSSC environmental genomes (clusters k, l, m, n, o, and p). The observed absence of rare clusters in either *Ralstonia* clade may be an artefact derived from biases in publicly available data.

We selected one member of each of the 16 *Ralstonia* T4SS gene clusters to be reference clusters (**Figure 2**, Supplemental Table S2B). Reference clusters a-j are from RSSC phytopathogen genomes, and reference clusters k-p are from non-RSSC environmental genomes. The references include T4SS genes predicted by multiple tools including CONJscan/MacSyFinder, NCBI CDD, and NCBI PGAP. The T4SS reference clusters contain 14-28 genes, with a median of 20.5 genes. The *virB4* genes range in size from 2,375 to 3,065 bp. To assign *vir/tra/trb/tfc* gene names to the reference clusters, we used CONJscan to classify the T4SS gene clusters into the established F/P/I/GI-type classification system. P-type systems use the *virB*/*D4* naming convention, GI-type systems use *tfc*, and F-type and I-type systems both use *tra*/*trb*. Notably, the F-type and I-type systems reuse certain gene names for non-homologous genes. Twelve clusters were classified as P-type T4SSs (clusters a, b, e, f, g, h, i, k, m, n, o, and p), two as F-type (clusters d and l), one as I-type (cluster j), and one as GI-type (cluster c) (**Figure 2**, Supplemental Figure S2).

**Figure 2.**
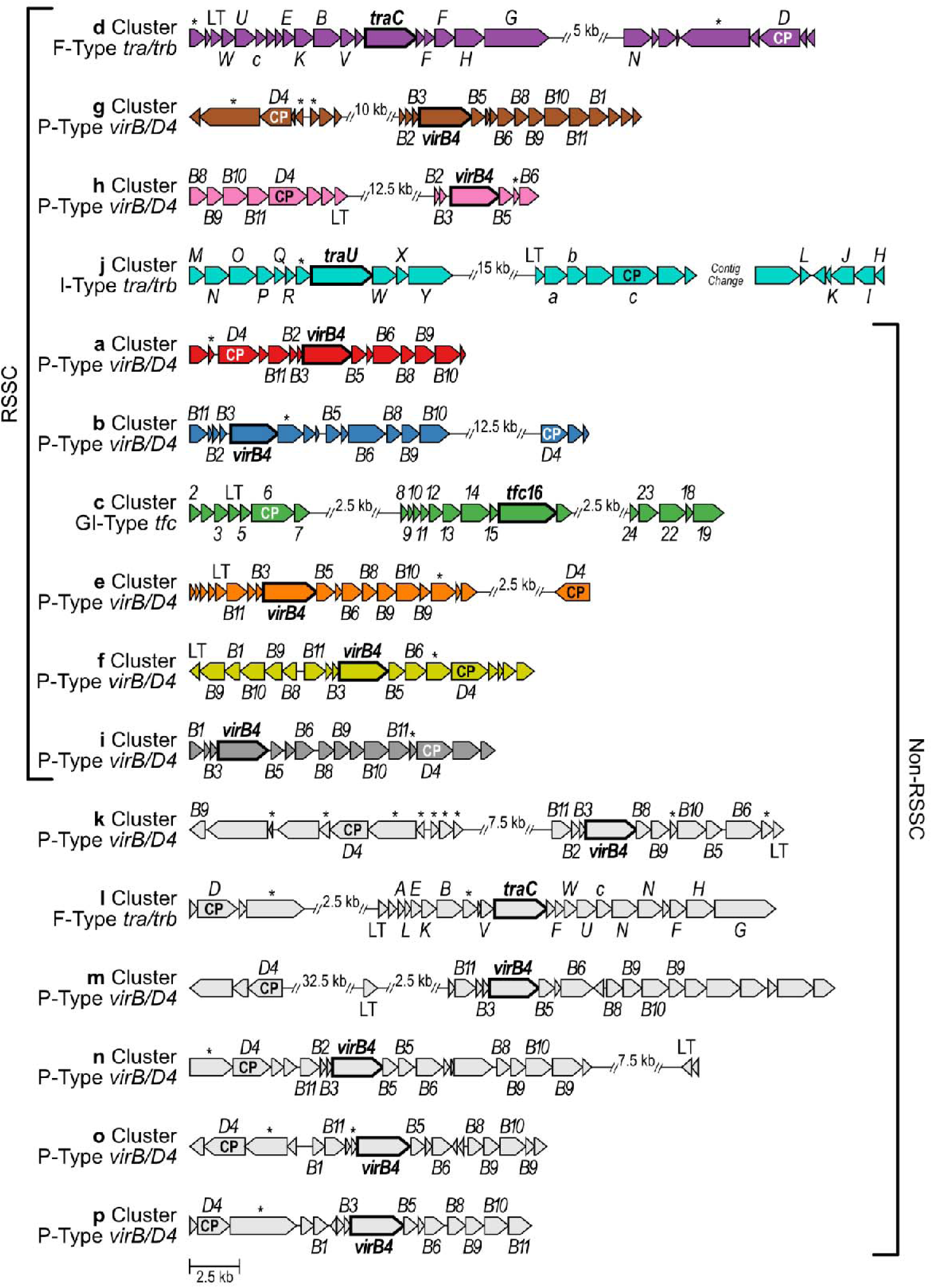
The T4SS gene clusters in *Ralstonia* have variable gene organization and composition. Clusters are ordered based on their phylogenetic distribution: found only in RSSC phytopathogens, found in both, or found only in non-RSSC environmentals. Genes in each cluster are labeled based on F/P/I/GI-type classification conventions: *tra*/*trb* (F-type), *virB*/*D4* (P-type), an independent *tra*/*trb* scheme (I-type), and *tfc* (GI-type). The *virB4* homologs have a bolded name (*traC*, *virB4*, *traU*, and *tfc16*) and a bolded gene arrow border. The genes annotated as the type IV coupling protein gene are labeled “CP”. “LT” indicates genes annotated as lytic transglycosylase genes. Genes marked with an asterisk (*) are T4SS genes that were not able to be annotated within the indicated F/P/I/GI-type convention. Gene clusters were defined using Operon-mapper, MacSyFinder v2 with CONJscan models, NCBI RefSeq annotations, and NCBI Conserved Domain Database. Gene clusters were visualized with clinker and labeled with Affinity Designer. Gaps are displayed where the representative cluster had regions without predicted T4SS genes; patterns of non-T4SS genes are not always consistent among all members of each cluster. The representative cluster j is found on two separate contigs in the only *Ralstonia* genome with cluster j. For additional details regarding genome, gene, and protein NCBI accessions, see Supplemental Table S2B and Supplemental Files S10-S26.

There are certain recurring gene organization patterns among the twelve P-type systems, despite the overall variability across the 16 T4SS gene clusters. These patterns in gene organization can be used to infer that a given T4SS gene cluster is a P-type T4SS. This is especially useful for more distantly related gene clusters where certain homologous genes have less than the 30% global amino acid identity, which is a default parameter for clinker. In all twelve cases, *virB4* is preceded by two small 225-417 bp genes on the same strand. In six cases, the two small genes are *virB2* and *virB3*, in that order. In the other six cases, only *virB3* is predicted by sequence alignment. The genes *virB8*, *virB9*, and *virB10* occur in succession, but in two clusters the *virB9* and *virB10* genes are separated by an additional gene. *virB11* either precedes *virB2*-*virB3*-*virB4* (8 clusters) or follows *virB8*-*virB9*-*virB10* (4 clusters). A second *virB9* paralog immediately follows *virB8*-*virB9*-*virB10* in four cases, only when that set of genes is not followed by *virB11*. To compare T4SSs in future genome sequences to the reference clusters, the GenBank files are available on Zenodo (DOI: 10.5281/zenodo.14873518; Supplemental Files S10-S26).

### T4SS gene clusters are more abundant in the non-RSSC environmentals than in the RSSC phytopathogens

Clinical pathogens are known to have abundant mobile genetic elements (1,2,6,55,56), and previous studies have established that RSSC phytopathogen genomes contain ICEs and genomic islands (57), transposable elements (58), and prophages (59). However, evolutionary comparisons of mobile genetic elements between pathogens and their relatives are less common than pathogen-centric analyses. Because RSSC phytopathogen genomes are known to be recombinogenic (60–62), both RSSC phytopathogen and non-RSSC environmental lineages are environmentally transmitted, and RSSC phytopathogen genomes are marginally larger than non-RSSC environmental genomes (RSSC median genome size, n=518: 5.6 Mb; non-RSSC median genome size, n=138: 5.4 Mb; Mann-Whitney test, *P* < 0.0001) (Supplemental Table S1), we hypothesized that T4SS gene clusters would be more abundant in the RSSC phytopathogens than in the non-RSSC environmentals.

We first looked at T4SS abundance qualitatively by visualizing presence and absence patterns on species trees of 636 RSSC phytopathogen genomes and 143 non-RSSC environmental genomes (**Figure 3A-B**). We queried the genomes for the VirB4 sequences from each reference cluster using multiple BLASTp searches, identifying 496 VirB4 sequences. The BLASTp results were consistent with the previous HMM search that led to Figure 1. Queries with VirB3 and VirB8 yielded a similar number of hits as VirB4 and protein trees with similar topologies (data not shown), but VirB3 and VirB8 searches only identified the P-type clusters. Nevertheless, during data curation, we identified and added five clusters with a putatively pseudogenized *virB4* gene to the tree (four cluster a and one cluster f). Overall, the T4SS gene clusters had a sporadic phylogenetic distribution across both the RSSC phytopathogens and the non-RSSC environmentals.

**Figure 3.**
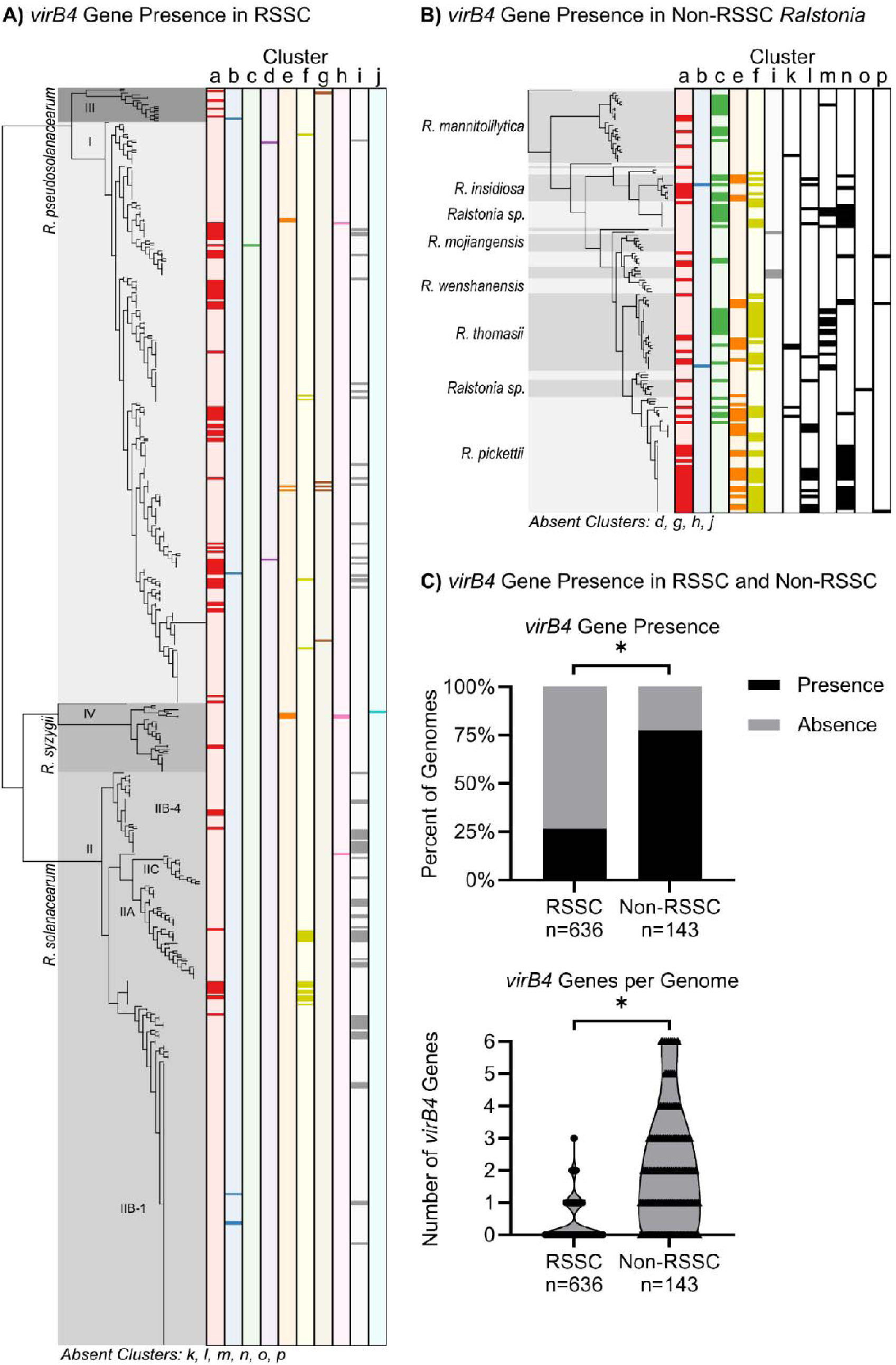
RSSC phytopathogen and non-RSSC environmentals had different abundance and phylogenetic distributions of sixteen T4SS clusters. Phylogenetic distribution of **A)** T4SS clusters a-j across 636 RSSC phytopathogen genomes and **B)** clusters a-c, e-f, i, and k-p across 143 non-RSSC environmental genomes. The species trees are approximately-maximum-likelihood trees built with FastTree 2 using 49 conserved protein sequences. The presence of the T4SS gene clusters was determined by BLASTp queries with each cluster’s representative VirB4 sequence, and synteny was manually checked for multiple hits per cluster. Each column represents one T4SS gene cluster, with the dark rectangles indicating presence of the cluster and the light background indicating absence of the cluster in each genome. For the non-RSSC environmental species tree, species are designated by alternating gray shading, with names labeled for large groups. The missing labels are, from top to bottom: *Ralstonia sp.*, *R. condita*, *Ralstonia sp.*, *R. flaminis*, *Ralstonia sp.*, *R. psammae*, *R. holmesii*, *R. chuxiongensis*, and *R. edaphis*. The trees with presence/absence data were visualized in iTOL. Supplemental Table S2A contains a list of *virB4* genes, and Supplemental Files S2 and S3 contain PDFs of the phylogenetic trees with labeled branches. **C)** Top: the proportion of RSSC phytopathogen and non-RSSC environmental genomes containing at least one T4SS marker gene (*virB4*). Fewer RSSC phytopathogen genomes contained *virB4* genes (Chi-Square test, * corresponds to ⍺ = 0.001). Bottom: the number of *virB4* copies per genome. Each point represents one genome. RSSC phytopathogen genomes had fewer T4SS gene clusters per genome than non-RSSC environmental genomes (Kolmogorov-Smirnov test, * corresponds to *P* < 0.0001).

Because the non-RSSC environmental genomes appeared to harbor more T4SS clusters, we quantified the abundance of T4SSs in each *Ralstonia* clade using the presence of *virB4* genes. We first quantified the incidence of any T4SS cluster, revealing the non-RSSC environmentals encoded more T4SS gene clusters (Chi-Square test, ⍺ = 0.001) (**Figure 3C**). Specifically, at least one T4SS gene cluster was found in 77.6% (111/143) of non-RSSC environmental genomes and 26.6% (169/636) of RSSC phytopathogen genomes. As a second metric, we quantified the number of T4SS gene clusters per genome using the presence of *virB4* genes. Once again, the non-RSSC environmental genomes contained more T4SS gene clusters (Kolmogorov-Smirnov test, *P* < 0.0001) (**Figure 3C**). The median number of T4SS gene clusters was 0 (IQR 0-1) in RSSC phytopathogen genomes and 2 (IQR 1-3) in non-RSSC environmental genomes. Of the genomes searched, no RSSC phytopathogen genomes had more than three T4SS gene clusters, while the non-RSSC environmental genomes contained up to six T4SS gene clusters.

We investigated whether the specific T4SS clusters varied in prevalence among RSSC phytopathogens and non-RSSC environmentals. When focusing only on the six T4SS gene clusters that were found in both the RSSC phytopathogens and non-RSSC environmentals (clusters a, b, c, e, f, and i), cluster a was consistently found in high prevalence and cluster b was consistently found in low prevalence. Cluster i was more common in the RSSC phytopathogens while clusters c, e, and f were more common in the non-RSSC environmentals.

### The RSSC phytopathogen genes RSp0179 and RSp1521 are unlikely to be T4SS-associated

Two genes (RSp0179/RS_RS18010 and RSp1521/RS_RS24455) in RSSC phytopathogen genomes have been described in the literature as T4SS genes (63,64) because they have homology to VirB10 and AcvB/VirJ-superfamilies. To determine whether RSp0179 and RSp1521 are T4SS-associated, we compared their homology to canonical VirB10 or AcvB/VirJ proteins, investigated their gene neighborhoods, and tested whether they share phylogenetic distributions with T4SS clusters.

VirB10 is a component of the transmembrane channel of the T4SS complex for P-type systems (10,11,15,16,65). We compared the sequences of RSp0179 to canonical VirB10 proteins from seven P-type T4SS clusters found in RSSC phytopathogen genomes (clusters a, b, e, f, g, h, and i). RSp0179 has an N-terminal signal peptide and a median length of 155 aa (range: 154-159 aa). In contrast, canonical VirB10 proteins are larger with median lengths of 414 aa (range: 404-493 aa). The short RSp0179 protein has homology with the middle of the VirB10 proteins; residues 8-135 of RSp0179 have 32% identity and 48% similarity to residues 174-326 in the RS_RS12910 protein (canonical VirB10 from cluster a in GMI1000). NCBI CDD annotates ∼200-400 aa of canonical VirB10 proteins with a VirB10/COG2948 domain, while the RSp0179 has ∼60 aa annotated as VirB10-like. Inspection of complete genomes showed that RSp0179 is located in a consistent location on the megaplasmid without any T4SS gene clusters (Figure S3A). To test whether RSp0179 has a sporadic phylogenetic distribution similar to the 16 T4SS clusters, we conducted BLASTp searches against the *Ralstonia* genus using thresholds of 80% identity and 80% coverage. RSp0179 was present in 94.9% (374/394) of RSSC phytopathogen genomes and none of the 143 non-RSSC environmental genomes. Because T4SSs are both sporadically present across the genus and less common in RSSC genomes than non-RSSC environmental genomes, the phylogenetic pattern of the VirB10-like protein RSp0179 does not correlate with T4SS clusters. Nevertheless, functional testing is required to understand the biological role of RSp0179.

Although AcvB/VirJ superfamily proteins are required for T4SS activity in some bacteria, experts have cautioned that the interaction may be indirect (66). Indeed, AcvB indirectly impacts *Agrobacterium* T4SS activity because AcvB is a periplasmic enzyme that directly affects inner membrane composition (67). *acvB* is encoded in an operon with *lpiA*. LpiA and AcvB contribute to lipid homeostasis by adding and removing lysine moieties to phosphatidylglycerol (PG) phospholipids (67). In *acvB* mutants, the elevated Lys-PG levels cause indirect inhibition of T4SS activity (67). Because RSp1521 is encoded in an *acvB*-like operon with a *lpiA* homolog (59.4% global amino acid similarity between CVI17646.1 and WP_011004769.1), we concur with a previous inference that RSp1521 is an *acvB* ortholog (68). Consistent with a role in lipid homeostasis, RSSC mutants lacking RSp1521 have pleiotropic phenotypes (68). Moreover, RSp1521 and model AcvB/VirJ-family proteins have the conserved Ser and His residues that are necessary for the Lys-PG hydrolase activity in the C-terminal domain (Supplemental File S37) (67,69). RSp1521/*acvB* is widely conserved in the *Ralstonia* genus. BLASTp searches with 82.8% identity and 78.5% coverage thresholds identified single-copy hits in 100% (143/143) of non-RSSC environmental genomes and 95.2% (375/394) of RSSC phytopathogen genomes. We also inspected the gene neighborhood of RSp1521/*acvB* and the *lpiA* homolog in complete genomes with and without T4SS gene clusters (Figure S3B). The putative operon was located in a consistent neighborhood on the megaplasmid and was not near any T4SS clusters, further supporting the hypothesis that RSp1521/*acvB* is not T4SS-associated.

### The T4SS gene clusters vary in prevalence and co-occurrence patterns in *Ralstonia* genomes

Since one genome can contain more than one T4SS gene cluster, we investigated which specific clusters co-occurred and in what quantities (**Figure 4A**). Out of the genomes that encoded at least one T4SS gene cluster, co-occurrence existed in 21.3% (36/169) of RSSC phytopathogen genomes and 71.2% (79/111) of non-RSSC environmental genomes. We found that co-occurrence was generally more common for more prevalent T4SS clusters (RSSC: clusters a, i, and f; non-RSSC: clusters a, f, and c). However, there were exceptions where low prevalence clusters co-occurred. Although RSSC phytopathogen cluster e occurred in only seven genomes, cluster e co-occurred twice each with other rare clusters g and h, which were only found in five and four genomes, respectively. Additionally, there were exceptions where two reasonably abundant clusters did not co-occur with each other; the non-RSSC environmentals contained 72 cluster a and 17 cluster m T4SSs, but no genome contained both. Both of these clusters commonly co-occurred with other T4SS clusters.

**Figure 4.**
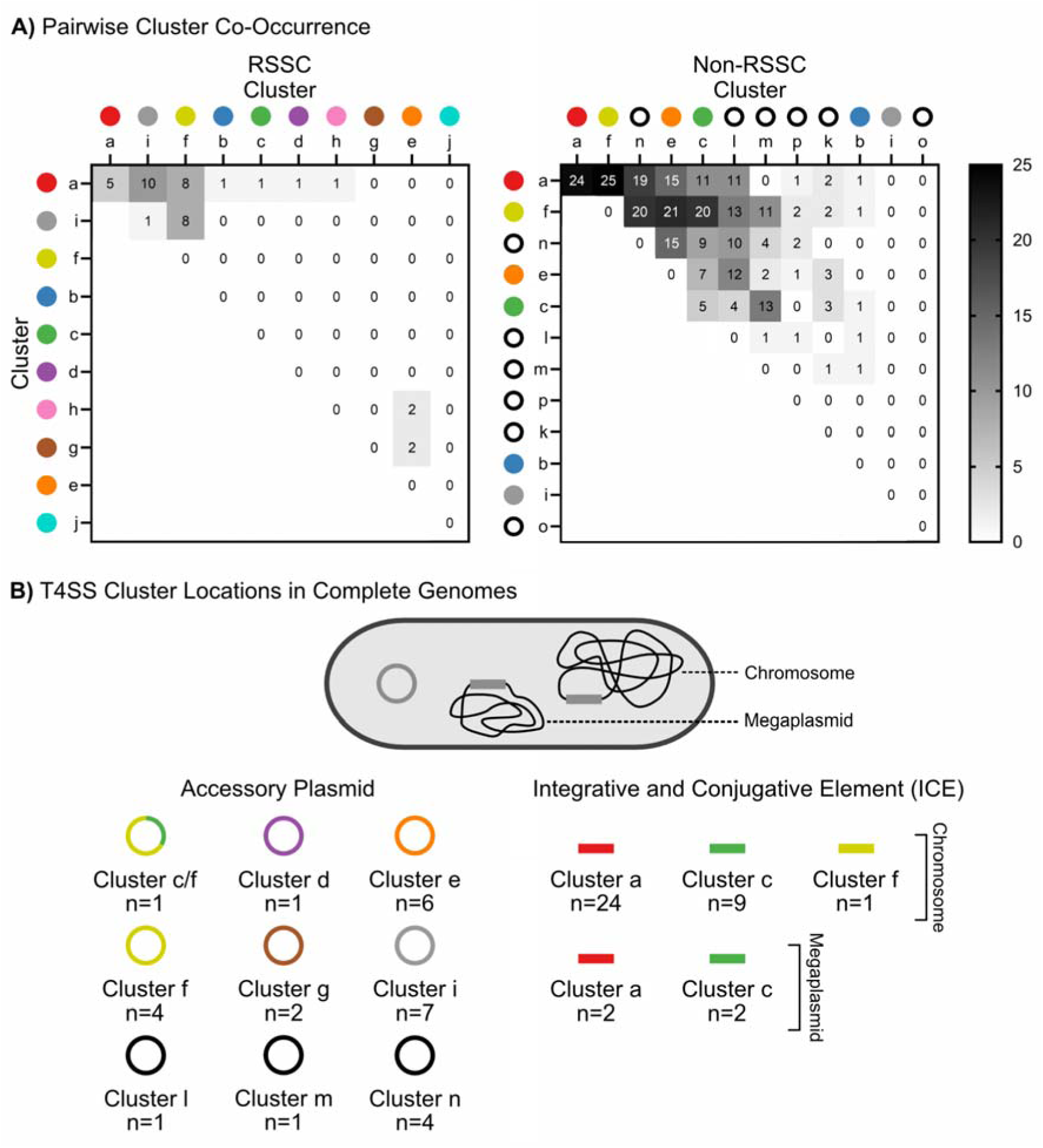
Genomic context sheds light on the ICE vs. accessory plasmid status of T4SS gene clusters. **A)** Pairwise comparison of T4SS cluster co-occurrence in genomes with two or more clusters. Numbers show how many genomes contain the indicated pair of T4SS gene clusters; genomes with more than two clusters are counted for each pairwise co-occurrence. Colored circles indicate clusters that were found in RSSC phytopathogen genomes, and black rings indicate clusters that were not found in the RSSC phytopathogens. **B)** T4SS gene clusters are encoded on accessory plasmids and ICEs. Rings indicate an accessory plasmid and rectangles indicate an ICE. T4SS cluster locations were determined using only complete genomes. Supplemental Table S2C shows the genomes used and the locus information for each T4SS region.

Although most cases of co-occurrence were between different cluster types, there were cases where two members of the same cluster exhibited co-occurrence (**Figure 4A**). Two or more cluster a T4SSs were present in five RSSC phytopathogen genomes and 24 non-RSSC environmental genomes. Additionally, two cluster c T4SSs were present in five non-RSSC environmental genomes. We ruled out a putative co-occurrence of cluster i in the draft genome of RSSC phytopathogen strain CIP221_UW464, because both occurrences were on opposite ends of the same contig. Those two putative cluster i copies had 99.95% nucleotide identity, so the dual presence likely reflects error assembling a ∼125 kb plasmid. Other than these instances, no other T4SS gene clusters were observed to co-occur with themselves in the same genome.

### The T4SS gene clusters are variably located on the chromosome, megaplasmid, and accessory plasmids

We investigated whether cluster a-p are predominantly maintained as accessory plasmids or ICEs in the bipartite genomes, which are composed of a ∼3.5 Mb chromosome and a ∼2.0 Mb megaplasmid. Assuming that members of the same cluster are likely to be part of the same plasmid incompatibility group (71), the co-occurrence data suggests the hypothesis that clusters a and c are not encoded on replicating plasmids. In 29 genomes, there were multiple cluster a T4SSs, and in five genomes there were two cluster c T4SSs (**Figure 4A**). If cluster a T4SSs were encoded on accessory plasmids, and all cluster a plasmids belonged to the same plasmid incompatibility group, we would not expect to find two different cluster a T4SSs in a single genome. However, the observed co-occurrences of distinct cluster a T4SSs in the same genome suggests that these clusters are encoded as ICEs.

To confidently determine the genomic location of the T4SS gene clusters, we narrowed our analysis to complete genomes: 33 T4SS gene clusters from 25 RSSC phytopathogen genomes and 33 T4SS gene clusters from 14 non-RSSC environmental genomes. With this dataset, we were able to infer locations of clusters a, c, d, e, f, g, i, l, m, and n, but there were no complete genomes containing clusters b, h, j, k, o, and p.

Analysis of complete genomes provided strong evidence that clusters a and c are often ICEs. We analyzed 26 cluster a T4SSs, which were encoded on the chromosome (n=24) or megaplasmid (n=2) (**Figure 4B**). For cluster c, we analyzed twelve T4SSs, revealing nine chromosomal clusters, two megaplasmid clusters, and one accessory plasmid cluster (**Figure 4B**). The cluster c on the accessory plasmid of non-RSSC environmental strain RRA could be an ICE that integrated into an existing plasmid because this plasmid contained a second T4SS (cluster f). Gonçalves *et al*. (2020) described six types of chromosomally-encoded ICEs that align with this paper’s cluster a (Tn*4371*, ICE*Rps1*, ICE*Rps2*, ICE*Rps3*, and ICE*Rsy1*) and cluster c (ICE*Rm1*) (57). Thus, we support the inference that clusters a and c are both generally maintained as ICEs.

The other clusters were predominantly present on accessory plasmids. For clusters d, e, g, i, l, m, and n, we analyzed one, six, two, seven, one, one, and four T4SS gene clusters, respectively. All of these clusters were identified on accessory plasmids (**Figure 4B**).

Moreover, the clusters did not confidently co-occur with variants of their own cluster type, further supporting their plasmid nature. For cluster f, we analyzed six T4SSs. We found one on a chromosome and five clusters on accessory plasmids, including the plasmid described above that also contained cluster c (**Figure 4B**). Since clusters f and c were identified on plasmids and as ICEs, it is possible that a cluster f ICE integrated into an existing cluster c plasmid. However, the high prevalence of cluster c in ICE format and cluster f in plasmid format suggests cluster c integrated into a cluster f plasmid. Analyzing a larger sample size of *Ralstonia* T4SSs can shed light on which are commonly maintained as accessory plasmids.

### At least fourteen T4SS gene clusters are predicted to be DNA conjugation systems

We hypothesized that the T4SS gene clusters in *Ralstonia* may encode self-transmissible DNA conjugation systems due to their sporadic distribution across the genus. Generally, the T4SSs that function as DNA conjugation systems require the proteins that facilitate the mobilization of the DNA cargo, the T4CP and relaxase. We searched for these mobilization genes in complete genomes containing T4SS gene clusters and also draft genomes if they contained a reference cluster from Figure 2, for a total of 38 T4SS gene clusters from 29 RSSC phytopathogen genomes and 39 T4SS gene clusters from 20 non-RSSC environmental genomes.

To identify T4CP genes, we first searched the 16 reference T4SS gene clusters using CONJscan/MacSyFinder. Then we used the synteny visualization tool clinker to confirm that the T4CPs were conserved in all 77 T4SS gene clusters searched. To search for relaxase genes in the vicinity of T4SS gene clusters, we used MOBscan. We identified a relaxase gene near 84.4% (65/77) of T4SS gene clusters searched. The T4SS gene clusters without an identifiable relaxase gene were cluster d (1/2), cluster e (7/7), and cluster m (2/2). Eight of these are clearly encoded on accessory plasmids, and the other two are reference clusters from draft genomes. Note that the absence of an identifiable relaxase gene is not sufficient to exclude DNA conjugation as a possible function. Overall, all 16 T4SS clusters appear to contain a T4CP (**Figure 2**) and 14 out of 16 are near a relaxase gene. Therefore, we predict that at least 14 gene clusters are DNA conjugation systems.

Because we used MOBscan to identify relaxase genes, we were able to determine the MOB class of each relaxase identified. MOBscan uses HMM profiles to classify MOB families. We found three of the twelve MOBscan MOB families in our dataset: P1, H, and F. A MOB_P1_ gene was associated with clusters a, b, h, i, j, and k; MOB_H_ with clusters c, f, n, and o; and MOB_F_ with clusters d, g, l, and p. In general, the MOB class of the relaxase gene near a T4SS gene cluster was consistent between members of the same cluster (Supplemental Table S2A). This consistency further supports the differentiation of clusters, such as in the case of the closely related clusters h (MOB_P1_) and p (MOB_F_).

Since we predicted that the majority of these T4SS gene clusters encode DNA conjugation systems, we investigated their putative regions of transfer. To do this, we analyzed 33 putative regions of transfer from 25 complete RSSC phytopathogen genomes and 33 putative regions of transfer from 14 complete non-RSSC environmental genomes. For accessory plasmids with a single T4SS cluster, the entire plasmid was considered the putative region of transfer. Otherwise, the putative region of transfer was determined by clinker synteny analysis, where the putative boundaries of the regions of transfer were inferred by comparing closely related genomes with and without T4SS clusters. On the plasmid with two T4SS clusters, synteny comparisons to single-T4SS plasmids and ICEs revealed the likely boundaries of clusters f and c, respectively. Collectively, the sizes of the putative regions of transfer were 32.6-389.8 kb (Figure S4A). Usually, members of the same cluster had similarly sized putative regions of transfer. For example, the regions for cluster a were 37.3-57.0 kb. In contrast, some regions had variable lengths, such as those from cluster f members where size ranged from 146.4 to 379.1 kb.

Additionally, we compared the GC contents of the putative regions of transfer and their parental genomes because differences in GC content could result from horizontal gene transfer (Figure S4B) (72). The putative regions in RSSC phytopathogen genomes had 57.2-64.3% GC, which was lower than the 66.5-67.0% GC of the whole genomes. For the non-RSSC environmentals, the regions had 58.2-67.0% GC compared to the 63.0-66.0% GC of the whole genomes. There was a significant difference in GC content between the regions and whole genomes for T4SS gene clusters in the RSSC phytopathogens (Wilcoxon test, *P* < 0.0001), but there was no significant difference in GC content for clusters in the non-RSSC environmentals (Wilcoxon test, *P* = 0.11). When considering GC content of the region vs. parental genome per cluster, the difference in GC content was significant for clusters a (Wilcoxon test, *P* < 0.0001), e (*P* = 0.031), f (*P* = 0.031), and i (*P* = 0.016), but not significant for clusters c (*P* = 0.30) and n (*P* = 0.13). Clusters d, g, l, and m had sample sizes too small for statistical comparison.

### RSSC phytopathogens and non-RSSC environmentals differ in cargo gene composition of T4SSs

Because the RSSC phytopathogens and non-RSSC environmentals have different lifestyles, we investigated if there was a difference in the cargo gene composition of their T4SSs’ putative regions of transfer. We focused on complete *Ralstonia* genomes to infer cargo gene functions.

The RSSC phytopathogens manipulate plants by using a type III secretion system (T3SS) to inject an arsenal of effector proteins into host cells (73), effector composition between RSSC phytopathogen genomes is highly variable (46), and effectors have been previously identified on an RSSC accessory plasmid (74). Thus, we hypothesized that RSSC phytopathogen T4SSs might contribute to the dynamic patterns by carrying T3 effectors and that T3 effectors would be more common in regions of transfer in RSSC phytopathogen T4SSs. Five of 33 RSSC phytopathogen genomes and zero of 33 non-RSSC environmental genomes had at least one T3 effector gene (**Figure 5**, Supplemental Table S2C), but there was not a significant difference in frequency (Mann-Whitney test, *P* = 0.053). The cluster i accessory plasmid in HA4-1 encoded one *ripP1* gene (also known as *ripBR* (74)) and two *ripAC* paralogs (also known as *ripBS* (74)). The cluster i plasmids in 068_2 and 077-080_1 each encoded a single copy of *ripBH*. The cluster i plasmid in RS24 and the cluster f plasmid in MolK2 each encoded a single copy of *ripBA*.

**Figure 5.**
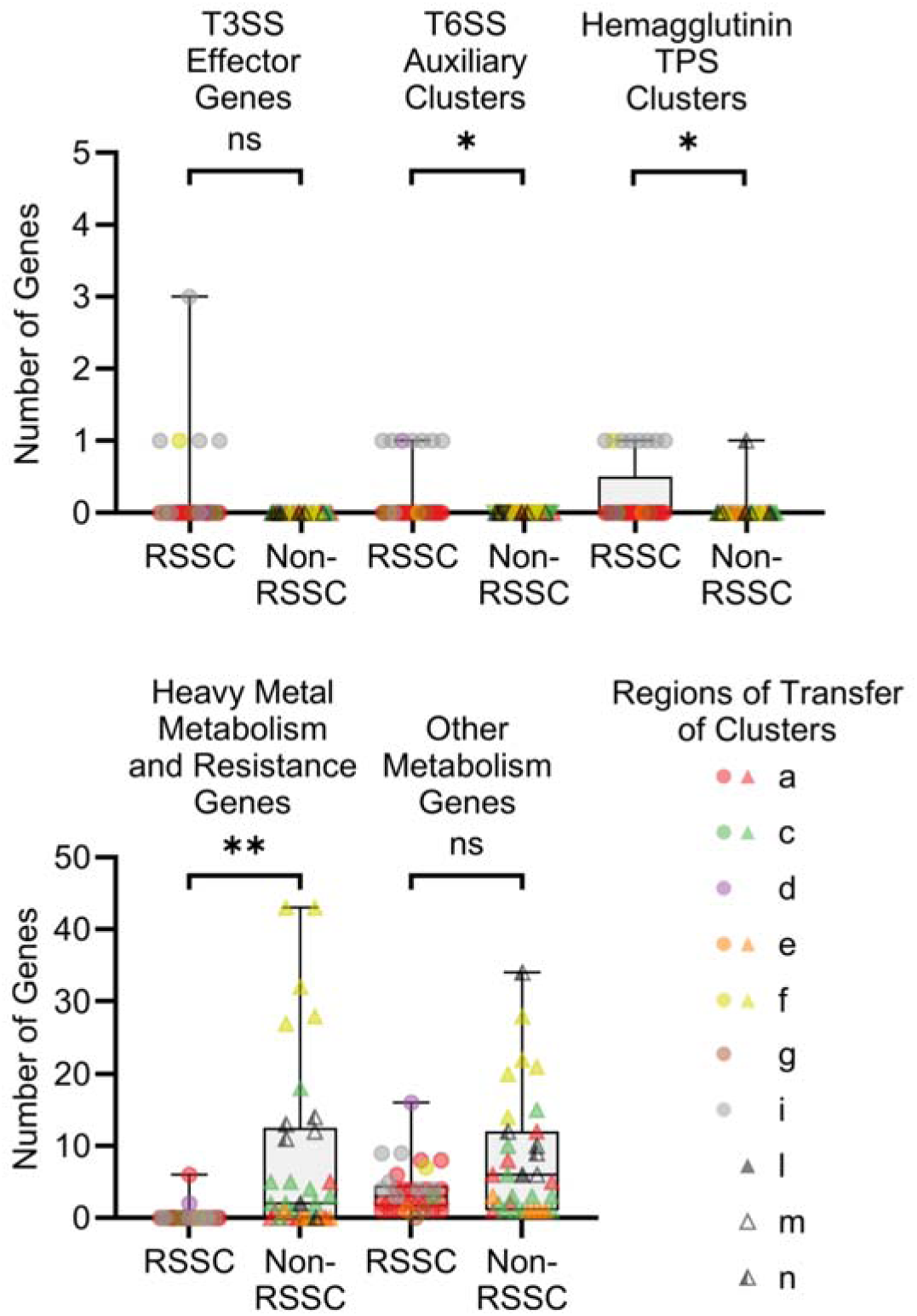
Cargo gene content of T4SS clusters differs between RSSC phytopathogens and non-RSSC environmentals. The graphs show the number of type III secretion system (T3SS) effector genes, type VI secretion system (T6SS) *vgrG*-linked auxiliary clusters, hemagglutinin two-partner secretion (TPS) clusters, heavy metal metabolism and resistance genes, and other metabolism genes present in each putative region of transfer. “Heavy metal” refers to transition (d-block) metals, post-transition (p-block) metals, and metalloids. Each region of transfer is represented by a circle (RSSC phytopathogen) or triangle (non-RSSC environmental) and is colored based on cluster identity. Significance was determined using the Mann-Whitney test: * corresponds to *P* < 0.05, ** corresponds to *P* < 0.0001, and ns corresponds to *P* > 0.05. Data was graphed and analyzed using GraphPad Prism v10.4.1. See Supplemental Table S2C for additional details about the specific genes identified in the putative regions of transfer. Supplemental Figure S4 conveys the GC content and sizes of the putative regions of transfer.

Recently, we showed that diverse classes of mobile genetic elements in RSSC phytopathogen genomes are often associated with microbial warfare-associated type VI secretion system (T6SS) genes, specifically the *vgrG*-linked auxiliary clusters with paired toxins and antitoxins (47). Several of the *vgrG*-linked toxin clusters were located on T4SS-containing plasmids, so we decided to investigate further. Here, we investigated the *Ralstonia* T4SS regions of transfer for the 25 previously classified *vgrG*-linked auxiliary clusters (47) and 19 additional *vgrG*-linked clusters from non-RSSC environmental genomes (Lowe-Power, unpublished). Consistent with the known enrichment of *vgrG*-linked auxiliary clusters in RSSC phytopathogen genomes (47), the RSSC phytopathogens had significantly more T6 *vgrG*-linked auxiliary clusters per region of transfer than the non-RSSC environmentals (Mann-Whitney test, *P* = 0.011) (**Figure 5**); seven of 33 RSSC phytopathogen genomes and zero of 33 non-RSSC environmental genomes had a single *vgrG*-linked auxiliary cluster (Supplemental Table S2C). The *vgrG*-linked auxiliary clusters were encoded on cluster i and d accessory plasmids of RSSC phytopathogens. In addition to the presence of *vgrG*-linked auxiliary clusters *aux6a* and *aux11* on four previously noted cluster i containing accessory plasmids (47), *aux11* was encoded on the cluster i plasmids in 068_2 and 077-080_1, and *aux6a* was present on the cluster d accessory plasmid of this study’s newly sequenced genome of RSSC strain Gazipur 4.

Filamentous hemagglutinin-associated two-partner secretion (TPS) system gene clusters have been previously identified in the RSSC phytopathogens (75) and hypothesized to function as contact-dependent inhibition systems participating in kin-recognition and competition (76,77). TPS systems are class b type V secretion systems (T5bSSs) known in other bacteria to be involved in diverse functions like surface adhesion, bacterial aggregation, contact-dependent inhibition, cytolysis/hemolysis, and iron acquisition (8,78–84). Hemagglutinin TPS clusters include a transporter gene (*fhaC*) and at least one hemagglutinin-repeat containing gene (*fhaB*/FHA) (78–84). The RSSC phytopathogens had significantly more hemagglutinin TPS clusters per region of transfer than the non-RSSC environmentals (Mann-Whitney test, *P* = 0.027) (**Figure 5**); eight of 33 RSSC phytopathogen genomes and one of 33 non-RSSC environmental genomes had a single hemagglutinin TPS cluster (Supplemental Table S2C). These clusters were encoded on seven cluster i accessory plasmids and one cluster f accessory plasmid in RSSC phytopathogen genomes. In the non-RSSC environmentals, there was one cluster n accessory plasmid in strain 12D that encoded a hemagglutinin TPS cluster.

Because the non-RSSC environmentals occupy more diverse environments than the RSSC phytopathogens, we reasoned that the non-RSSC environmentals might benefit from T4SS-associated metabolism genes. To identify metabolism genes in the complete genome T4SS regions of transfer, we used four tools: ModelSEED v2 (50), METGeneDb v2 (51), DRAM (48), and dbCAN2 (49). We separated the aggregated results into two categories: (i) heavy metal metabolism and resistance genes and (ii) other metabolism genes. Here, the term “heavy metal” refers to transition (d-block) metals, post-transition (p-block) metals, and metalloids. Other metabolism genes were defined as any metabolism genes other than those identified as heavy metal metabolism and resistance genes.

Heavy metal metabolism and resistance genes were found in 22/33 non-RSSC environmental regions of transfer but only 2/33 RSSC phytopathogen regions of transfer (Supplemental Table S2C). The non-RSSC environmentals had significantly more heavy metal metabolism and resistance genes per region of transfer than the RSSC phytopathogens (Mann-Whitney test, *P* < 0.0001) (**Figure 5**). Within the non-RSSC environmentals, the number of heavy metal metabolism and resistance genes per region of transfer was 1-43 in the 22 regions identified. These genes were identified in regions of transfer of diverse clusters: cluster a (n=2), c (n=9), e (n=1), f (n=5), l (n=1), m (n=1), and n (n=3). Gene annotations suggest that these genes are related to mercury, nickel, zinc, cadmium, lead, cobalt, arsenic, chromium, iron, copper, silver, antimony, and tellurium interactions. Within the RSSC phytopathogens, six heavy metal metabolism and resistance genes were identified within a cluster a region of transfer in strain UW386, and two genes were identified on a cluster d accessory plasmid in strain Gazipur 4. Gene annotations suggest that these genes are related to mercury and cobalt interactions, respectively.

Genes for other metabolic functions were found in 32/33 RSSC phytopathogen regions of transfer and 33/33 non-RSSC environmental regions of transfer (Supplemental Table S2C). Within the RSSC phytopathogens, the number of other metabolism genes per region of transfer was 1-16 in the 32 regions identified. The single RSSC phytopathogen region for which we did not identify other metabolism genes was the cluster g accessory plasmid in strain SL1931. Within the non-RSSC environmentals, the number of other metabolism genes per region of transfer was 1-34. Despite the trend, there was no significant difference in the number of other metabolism genes per region of transfer between the RSSC phytopathogens and the non-RSSC environmentals (Mann-Whitney test, *P* = 0.089) (**Figure 5**). An annotated clinker visualization of the cluster a putative regions of transfer is available in Supplemental Figure S5. Clinker files for putative regions of transfer from each of the other clusters with complete genomes are available in Supplemental Files S27-S36.

### Specialization of a T4SS cluster to the RSSC phytopathogens

We hypothesized that cluster i might have non-exclusively specialized to the RSSC phytopathogens based on the previous results. As described, cluster i T4SSs are more prevalent in RSSC phytopathogens (n=71) as compared to non-RSSC environmentals (n=4). This is in contrast to the overall trend of greater T4SS prevalence in the non-RSSC environmentals (**Figure 3**).

Closer analysis of the VirB4 phylogeny (**Figure 1**) revealed that the cluster i T4SSs in the RSSC phytopathogens were almost entirely grouped by the species of *Ralstonia* they came from (**Figure 6**). However, the branching pattern is not perfectly congruent with the species tree made from the genomes that the cluster i T4SSs were found in. In contrast, the VirB4 phylogenies for other prevalent T4SS gene clusters, such as cluster a, were not grouped by the *Ralstonia* species in which the VirB4 sequences were identified. The cluster i VirB4 phylogeny has four major clades (**Figure 6**). The first clade is the outgroup, which includes two VirB4 sequences from one phylotype II *R. solanacearum* genome (Rs545_UW752) and one *R. chuxiongensis* genome. The second clade includes six VirB4 sequences from three phylotype I *R. pseudosolanacearum* genomes, two *R. chuxiongensis* genomes, and one *R. psammae* genome. The third clade includes 16 VirB4 sequences from 16 phylotype I *R. pseudosolanacearum* genomes. The fourth clade includes 51 VirB4 sequences from 49 phylotype II *R. solanacearum* genomes and one phylotype I *R. pseudosolanacearum* genome (RALF-MA). The RSSC phytopathogen phylotypes exhibit different geographical distributions, with phylotype I *R. pseudosolanacearum* originating in Asia and phylotype II *R. solanacearum* originating in the Americas. The phylotype I *R. pseudosolanacearum* strain RALF-MA was isolated from Costa Rica, most likely having been introduced to the Americas in the last few hundred years. Therefore, the origin of the cluster i T4SS in RALF-MA could be a phylotype II *R. solanacearum* strain.

**Figure 6.**
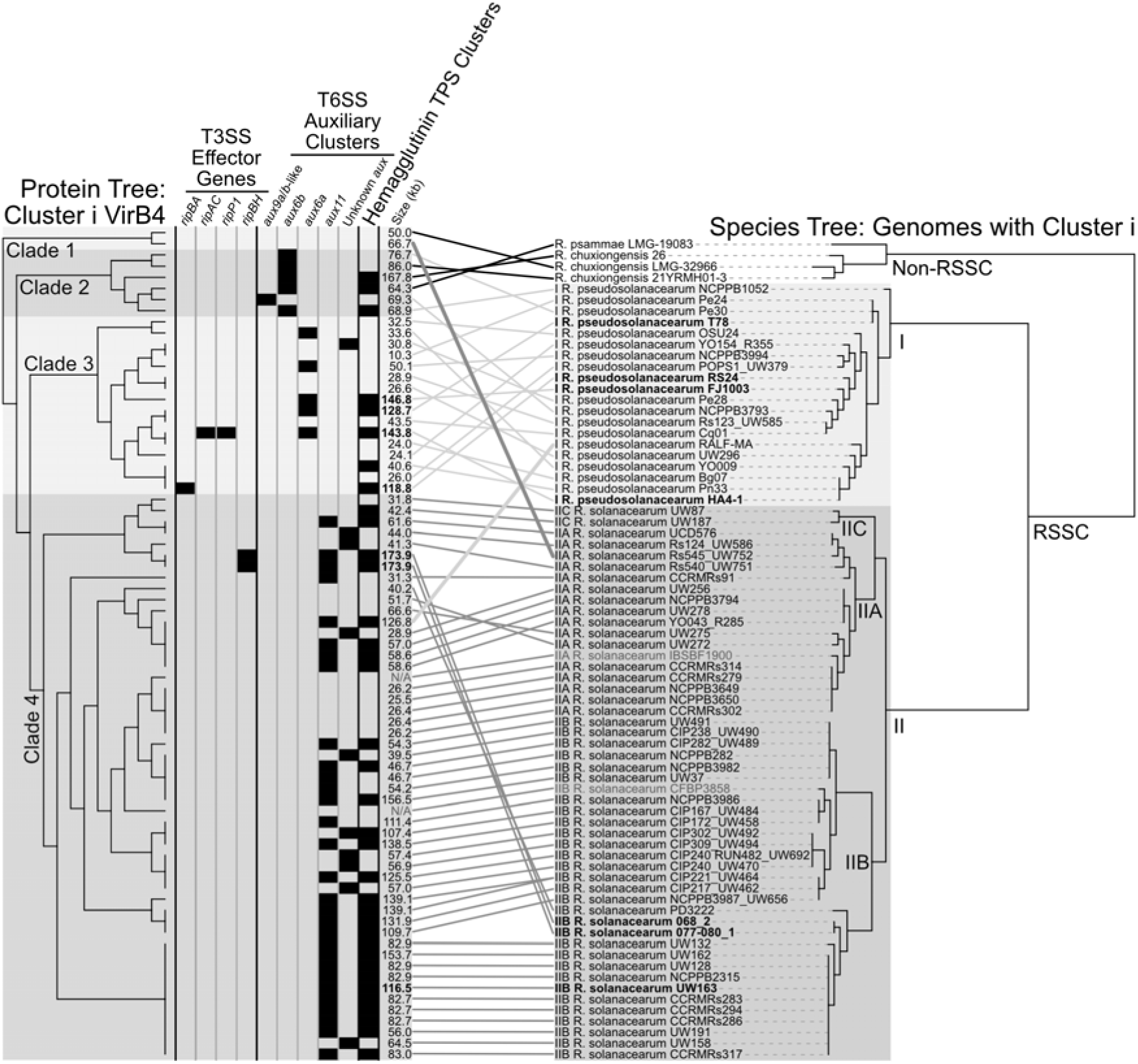
Phylogenetic analysis suggests cluster i T4SSs evolved a specialized relationship with RSSC phytopathogens. The left cladogram is a VirB4 protein tree made from an MSA of 75 cluster i sequences. The right tree is a species tree made from the 74 *Ralstonia* genomes that contain a cluster i T4SS. The columns show the presence of specific T3 effector genes, T6 *vgrG*-linked auxiliary clusters, and hemagglutinin TPS clusters in each cluster i T4SS. The sizes of the putative regions of transfer are noted, bold indicates the cluster is encoded in a complete genome, and grayed “N/A” labels are listed for T4SSs encoded in old genomes with contigs randomly concatenated into one scaffold. Branches of the phylogenetic trees were rotated manually in iTOL, and the tanglegram linkages and labels were added in Affinity Designer. See Supplemental File S38 for a cluster i region average nucleotide identity (ANI) matrix and Supplemental File S39 for a cluster i region clinker visualization.

We computed an ANI matrix for cluster i putative regions of transfer, which includes T4SS genes. Seven cluster i VirB4 sequences were found in complete genomes and 68 in draft genomes (n=75 total). For clusters in draft genomes, we treated the putative region of transfer as the whole contig that encoded the *virB4* gene. Two of the 75 cluster i T4SSs were found in nearly identical copies on either end of the same contig and were treated as one cluster. Additionally, two clusters were found in early genomes that had hundreds of contigs concatenated randomly into one sequence before submission to NCBI; these were excluded from the analysis because we could not confidently delineate contig boundaries. The ANI matrix was computed with ANIb using pyANI-plus for the 72 remaining regions. Hierarchical clustering using Morpheus resulted in a dendrogram with the same four major clades as the VirB4 phylogeny (Supplemental File S38).

Our cargo gene analysis revealed that the T3 effectors, T6 *vgrG*-linked auxiliary clusters, and hemagglutinin TPS clusters were common within cluster i regions (**Figure 5**). To discover if the trend was consistent, we searched for these putatively adaptive genes in 72 cluster i putative regions of transfer from draft and complete *Ralstonia* genomes (Supplemental Table S2D, Supplemental File S39). Four regions (5.6%) had at least one T3 effector gene and these regions were all previously described accessory plasmids from complete genomes (**Figures 5** and **6**). These regions were grouped within clades 3 and 4 of the cluster i VirB4 phylogeny. We identified a single T6 *vgrG* gene in 50 regions (69.4%), five from *aux6a*, five from *aux6b*, 29 from *aux11*, and a unique *aux9*-like cluster with divergent gene content from the previously classified *aux9a* and *aux9b* (**Figure 6**) (47). Ten of the identified *vgrG* genes were not able to be matched with a particular auxiliary cluster because they were truncated at the ends of contigs. The different T6 auxiliary clusters corresponded with the clades of the VirB4 phylogeny: *aux6b* and *aux9a*/*b*-like in clade 2, *aux6a* in clade 3, and *aux11* in clade 4. The identified T6 auxiliary clusters were generally in similar locations a few genes downstream of the T4SS cluster. No *vgrG* genes were identified in the remaining 22 regions, however, at least 16 of those regions had contigs that ended before the expected location of a T6 auxiliary cluster. We identified hemagglutinin TPS clusters in 35 regions of transfer (48.6%) (**Figure 6**). Of these clusters, 17 were whole clusters, meaning they included both *fhaC* and *fhaB* genes. The 18 partial clusters had at least one component of a TPS cluster, with the other part truncated by the ends of contigs. These TPS clusters were found in clades 2, 3, and 4 of the VirB4 phylogeny.

## Discussion

Contrary to our initial hypothesis, T4SS were less prevalent in RSSC phytopathogen genomes than in non-RSSC environmental genomes. Notably, there were functional differences in the genetic cargo associated with the T4SSs. What does this mean about the eco-evolutionary differences between the three RSSC phytopathogen species and the 17 non-RSSC environmental species?

Assuming that this study’s T4SS clusters are *bona fide* conjugative systems, the non-RSSC environmental species may inhabit bacterial communities with a greater diversity of conjugative donors, have fewer physiological barriers to acquiring and maintaining conjugated DNA, or more often benefit from the functions conferred by the conjugative elements. The prevalence of heavy metal metabolism and resistance genes in non-RSSC environmental putative regions of transfer may reflect their broader environmental distribution, in which exposure to toxic heavy metals may be more frequent. Correspondingly, many bacteria have evolved various mechanisms to resist the toxicity (85,86) and/or metabolize these metals. Therefore, the enrichment of heavy metal resistance genes in the putative T4SS regions of transfer is consistent with horizontal acquisition of adaptive traits.

The three monophyletic RSSC phytopathogen species have a more tailored and predictable life cycle of plant infection and environmental survival in soil and surface water. The phytopathogens’ lower prevalence of T4SSs is surprising because they are known to be naturally transformable (60,87). Additionally, T4SSs benefit other phytopathogens by conferring virulence (88,89). Thus, the RSSC phytopathogens might encounter barriers to acquiring conjugated DNA. For example, they may encounter a limited pool of non-kin cells that could donate conjugative elements during their explosive replication in plants. Nevertheless, we and others have identified ecologically beneficial genetic cargo on some of the T4SS regions of transfer in RSSC phytopathogen genomes: T3 effectors, and auxiliary T6 toxin clusters and hemagglutinin TPS clusters (46,47,57,74–77). These three classes of genes mediate or putatively mediate interactions with RSSC phytopathogens’ ecological targets: plants (T3 effectors) and microbial competitors (T6 toxins and hemagglutinin TPS toxins). At the genome level, it has been previously established that T6 auxiliary clusters are enriched in the RSSC phytopathogens (47), and we speculate that systematic comparison will also show that phytopathogen genomes are also enriched in T3 effectors and hemagglutinin TPS clusters.

Interestingly, cluster i appears to have a preferential association with RSSC phytopathogens. Their regions of transfer often encoded T6 *vgrG*-linked auxiliary clusters and hemagglutinin TPS clusters, while they occasionally encoded T3 effector genes. Cluster i T4SSs were enriched in the RSSC phytopathogens as compared to the non-RSSC environmentals. Additionally, in the VirB4 phylogeny, the cluster i T4SSs in the RSSC phytopathogens were strongly clustered by the species of *Ralstonia* they came from. Based on these results, we speculate that cluster i T4SSs are adapted to RSSC phytopathogens.

Several of the T4SS gene clusters that we named have been previously identified in comparative genomics papers: cluster a ICEs (57,63,74,75,90–93), cluster c ICEs (57), cluster g plasmids (63,74,90,94), and cluster i plasmids (74,91). Additionally, Morales and Sequeira (1985) identified and estimated the sizes of several accessory plasmids in RSSC phytopathogen strains, including a 120 MDa (∼197 kb) plasmid in UW256, which likely encodes the cluster i T4SS identified in the UW256 genome (95). Although multiple T4SS clusters are present on accessory plasmids, there are also accessory plasmids in *Ralstonia* genomes that lack T4SS gene clusters (data not shown). One of these T4SS-null accessory plasmids co-occurs with a cluster i plasmid in UW163, so it is possible that it is mobilizable by the cluster i T4SS.

This study has several limitations. As with any survey of public genomic data, there is an inherent sampling bias in publicly available data, and our dataset does not capture the total biological variation of this genus. For example, the observation that certain clusters were not present in the available genomes could be an artefact of the analyzed dataset. Although more diverse *Ralstonia* genomes have become available in recent years, the genomes of environmental *Ralstonia* were underrepresented in our dataset by 4.5-fold. Additionally, our T4SS search approach likely yielded occasional false negatives against the analyzed genomes because the HMMER and BLASTp searches queried translated amino acid sequences. We partially mitigated this risk by manually curating five additional T4SS gene clusters with putatively pseudogenized *virB4* genes that were embedded within otherwise intact T4SS loci. These clusters were retained because their genomic context indicates that the T4SS was acquired by an ancestor of the genome-sequenced strain, even if loss-of-function mutations may have arisen later. False negatives could also have occurred in draft genomes due to incomplete sequencing and/or assembly of the T4SS clusters despite high CheckM completeness statistics. Another limitation lies in the identification of metabolism-related genetic cargo. Although we used four databases to identify metabolic genes, there are certainly metabolic proteins that are unrepresented in available databases.

We used the identification of a T4CP gene and a relaxase gene to infer that most of the identified T4SSs could be DNA conjugation systems. This inferential approach does not predict whether a specific T4SS may be operational *in vivo*. It also cannot differentiate between DNA conjugation and DNA release systems because experimental testing is needed, and we suggest conjugation assays as a method of experimental validation in the future. However, DNA release systems have only rarely been identified; DNA release systems are limited to the T4SS encoded on the gonococcal genetic island (GGI) in *Neisseria gonorrhoeae* (96,97) and a T4SS in nontypeable *Haemophilus influenzae* (98). Therefore, it is more likely that a T4SS with a T4CP and an associated relaxase functions as a DNA conjugation system rather than a DNA release system. Although, the possibility that these T4SSs could function as DNA release systems introduces an alternative hypothesis for their mechanism of transmission to *Ralstonia* species. It has been determined that RSSC phytopathogens are naturally transformable in multiple conditions including *in vitro* and *in planta*, and are able to take up exogenous DNA ranging in size from 30 to 90 kb (60,87). Therefore, natural transformation has the potential to contribute to T4SS acquisition. DNA release via a T4SS could provide the exogenous DNA for natural transformation. This is the mechanism of horizontal transfer of the GGI of *Neisseria gonorrhoeae* (96,97). The GGI is released by its T4SS, then the naturally transformable *Neisseria* spp. in the vicinity recognize a specific sequence in the released DNA that triggers uptake by natural transformation (96,99). Nevertheless, the circumstantial evidence and Occam’s razor suggest most of the T4SSs in this study are part of self-transmissible mobile genetic elements.

Overall, the differences in T4SS prevalence and cargo gene composition between the RSSC phytopathogens and non-RSSC environmentals likely reflect differences in their ecological niches and lifestyles (100).

## Author contributions

TCC: Conceptualization, Data curation, Analysis, Experiments / Investigation, Supervision, Visualization, Writing – original draft, MLCA: Conceptualization, Project administration, Supervision, Writing – review & editing, GYS: Conceptualization, Experiments / Investigation, Writing – review & editing, AJB: Resources, Supervision, Writing – review & editing, SCDC: Experiments / Investigation, Resources, Writing – review & editing, KN: Resources, Writing – review & editing, AS: Resources, Writing – review & editing, TMLP: Conceptualization, Project administration, Resources, Supervision, Writing – review & editing.

## Acknowledgements

We would like to acknowledge Titus Brown and Clay Fuqua for their valuable discussions, and Jonathan Jacobs and Taylor Klass for sharing unpublished genomes.

## References

1. Hazen TH, Pan L, Gu JD, Sobecky PA. The contribution of mobile genetic elements to the evolution and ecology of *Vibrios*. FEMS Microbiology Ecology. 2010;74(3): 485–499. 10.1111/j.1574-6941.2010.00937.x.

2. Tenaillon O, Skurnik D, Picard B, Denamur E. The population genetics of commensal *Escherichia coli*. Nature Reviews Microbiology. 2010;8(3): 207–217. 10.1038/nrmicro2298.

3. Savory EA, Fuller SL, Weisberg AJ, Thomas WJ, Gordon MI, Stevens DM, et al. Evolutionary transitions between beneficial and phytopathogenic *Rhodococcus* challenge disease management. eLife. 2017;6: e30925. 10.7554/eLife.30925.

4. Martínez JL. Ecology and Evolution of Chromosomal Gene Transfer between Environmental Microorganisms and Pathogens. Baquero F, Bouza E, Gutiérrez-Fuentes JA, Coque TM (eds) Microbiology Spectrum. 2018;6(1). 10.1128/microbiolspec.MTBP-0006-2016.

5. Melnyk RA, Hossain SS, Haney CH. Convergent gain and loss of genomic islands drive lifestyle changes in plant-associated *Pseudomonas*. The ISME Journal. 2019;13(6): 1575–1588. 10.1038/s41396-019-0372-5.

6. Gyles C, Boerlin P. Horizontally Transferred Genetic Elements and Their Role in Pathogenesis of Bacterial Disease. Veterinary Pathology. 2014;51(2): 328–340. 10.1177/0300985813511131.

7. Fernandes AS, Gonçalves OS, De Lima LMO, Santana MF. Does the mobilome of *Ralstonia solanacearum* influence the evolution and virulence of this pathogen? Tropical Plant Pathology. 2025;50(1): 29. 10.1007/s40858-025-00718-z.

8. Chang JH, Desveaux D, Creason AL. The ABCs and 123s of Bacterial Secretion Systems in Plant Pathogenesis. Annual Review of Phytopathology. 2014;52(1): 317–345. 10.1146/annurev-phyto-011014-015624.

9. Johnson CM, Grossman AD. Integrative and Conjugative Elements (ICEs): What They Do and How They Work. Annual Review of Genetics. 2015;49(1): 577–601. 10.1146/annurev-genet-112414-055018.

10. Alvarez-Martinez CE, Christie PJ. Biological Diversity of Prokaryotic Type IV Secretion Systems. Microbiology and Molecular Biology Reviews. 2009;73(4): 775–808. 10.1128/MMBR.00023-09.

11. Juhas M, Crook DW, Hood DW. Type IV secretion systems: tools of bacterial horizontal gene transfer and virulence. Cellular Microbiology. 2008;10(12): 2377–2386. 10.1111/j.1462-5822.2008.01187.x.

12. Fronzes R, Christie PJ, Waksman G. The structural biology of type IV secretion systems. Nature Reviews Microbiology. 2009;7(10): 703–714. 10.1038/nrmicro2218.

13. Guglielmini J, De La Cruz F, Rocha EPC. Evolution of Conjugation and Type IV Secretion Systems. Molecular Biology and Evolution. 2013;30(2): 315–331. 10.1093/molbev/mss221.

14. Costa TRD, Patkowski JB, Macé K, Christie PJ, Waksman G. Structural and functional diversity of type IV secretion systems. Nature Reviews Microbiology. 2024;22(3): 170–185. 10.1038/s41579-023-00974-3.

15. Christie PJ. The Mosaic Type IV Secretion Systems. Lovett ST, Bernstein HD (eds) EcoSal Plus. 2016;7(1). 10.1128/ecosalplus.esp-0020-2015.

16. Cascales E, Christie PJ. The versatile bacterial type IV secretion systems. Nature Reviews Microbiology. 2003;1(2): 137–149. 10.1038/nrmicro753.

17. Garcillán-Barcia MP, Francia MV, De La Cruz F. The diversity of conjugative relaxases and its application in plasmid classification. FEMS Microbiology Reviews. 2009;33(3): 657–687. 10.1111/j.1574-6976.2009.00168.x.

18. Prior P, Ailloud F, Dalsing BL, Remenant B, Sanchez B, Allen C. Genomic and proteomic evidence supporting the division of the plant pathogen *Ralstonia solanacearum* into three species. BMC Genomics. 2016;17(1): 90. 10.1186/s12864-016-2413-z.

19. Safni I, Subandiyah S, Fegan M. Ecology, Epidemiology and Disease Management of *Ralstonia syzygii* in Indonesia. Frontiers in Microbiology. 2018;9: 419. 10.3389/fmicb.2018.00419.

20. Ryan MP, Adley CC. *Ralstonia* spp.: emerging global opportunistic pathogens. European Journal of Clinical Microbiology & Infectious Diseases. 2014;33(3): 291–304. 10.1007/s10096-013-1975-9.

21. Bartoli C, Roux F, Lamichhane JR. Molecular mechanisms underlying the emergence of bacterial pathogens: an ecological perspective. Molecular Plant Pathology. 2016;17(2): 303–310. 10.1111/mpp.12284.

22. Parks DH, Imelfort M, Skennerton CT, Hugenholtz P, Tyson GW. CheckM: assessing the quality of microbial genomes recovered from isolates, single cells, and metagenomes. Genome Research. 2015;25: 1043–1055. 10.1101/gr.186072.114.

23. Arkin AP, Cottingham RW, Henry CS, Harris NL, Stevens RL, Maslov S, et al. KBase: The United States Department of Energy Systems Biology Knowledgebase. Nature Biotechnology. 2018;36(7): 566–569. 10.1038/nbt.4163.

24. Greenrod STE, Stoycheva M, Elphinstone J, Friman VP. Global diversity and distribution of prophages are lineage-specific within the *Ralstonia solanacearum* species complex. BMC Genomics. 2022;23: 689. 10.1186/s12864-022-08909-7.

25. Bolger AM, Lohse M, Usadel B. Trimmomatic: a flexible trimmer for Illumina sequence data. Birol I (ed.) Bioinformatics. 2014;30(14): 2114–2120. 10.1093/bioinformatics/btu170.

26. Wick RR. Filtlong. GitHub. https://github.com/rrwick/Filtlong

27. Bankevich A, Nurk S, Antipov D, Gurevich AA, Dvorkin M, Kulikov AS, et al. SPAdes: A New Genome Assembly Algorithm and Its Applications to Single-Cell Sequencing. Journal of Computational Biology. 2012;19(5): 455–477. 10.1089/cmb.2012.0021.

28. Wick RR, Judd LM, Gorrie CL, Holt KE. Unicycler: Resolving bacterial genome assemblies from short and long sequencing reads. PLOS Computational Biology. 2017;13(6): e1005595. 10.1371/journal.pcbi.1005595.

29. Seemann T. Prokka: rapid prokaryotic genome annotation. Valencia A (ed.) Bioinformatics. 2014;30(14): 2068–2069. 10.1093/bioinformatics/btu153.

30. Elphinstone JG, Hennessy J, Wilson JK, Stead DE. Sensitivity of different methods for the detection of *Ralstonia solanacearum* in potato tuber extracts. EPPO Bulletin. 1996;26(3–4): 663–678. 10.1111/j.1365-2338.1996.tb01511.x.

31. Kelman A. The relationship of pathogenicity of *Pseudomonas solanacearum* to colony appearance in a tetrazolium medium. Phytopathology. 1954;44(12): 693–695.

32. Kolmogorov M, Yuan J, Lin Y, Pevzner PA. Assembly of long, error-prone reads using repeat graphs. Nature Biotechnology. 2019;37(5): 540–546. 10.1038/s41587-019-0072-8.

33. Gilchrist CLM, Chooi YH. clinker & clustermap.js: automatic generation of gene cluster comparison figures. Robinson P (ed.) Bioinformatics. 2021;37(16): 2473–2475. 10.1093/bioinformatics/btab007.

34. Camacho C, Coulouris G, Avagyan V, Ma N, Papadopoulos J, Bealer K, et al. BLAST+: architecture and applications. BMC Bioinformatics. 2009;10(1): 421. 10.1186/1471-2105-10-421.

35. Edgar RC. MUSCLE: multiple sequence alignment with high accuracy and high throughput. Nucleic Acids Research. 2004;32(5): 1792–1797. 10.1093/nar/gkh340.

36. Eddy SR. Accelerated Profile HMM Searches. Pearson WR (ed.) PLoS Computational Biology. 2011;7(10): e1002195. 10.1371/journal.pcbi.1002195.

37. Letunic I, Bork P. Interactive Tree of Life (iTOL) v6: recent updates to the phylogenetic tree display and annotation tool. Nucleic Acids Research. 2024;52(W1): W78–W82. 10.1093/nar/gkae268.

38. Castresana J. Selection of Conserved Blocks from Multiple Alignments for Their Use in Phylogenetic Analysis. Molecular Biology and Evolution. 2000;17(4): 540–552. 10.1093/oxfordjournals.molbev.a026334.

39. Price MN, Dehal PS, Arkin AP. FastTree 2 – Approximately Maximum-Likelihood Trees for Large Alignments. Poon AFY (ed.) PLoS ONE. 2010;5(3): e9490. 10.1371/journal.pone.0009490.

40. Tatusova T, DiCuccio M, Badretdin A, Chetvernin V, Nawrocki EP, Zaslavsky L, et al. NCBI prokaryotic genome annotation pipeline. Nucleic Acids Research. 2016;44(14): 6614–6624. 10.1093/nar/gkw569.

41. Wang J, Chitsaz F, Derbyshire MK, Gonzales NR, Gwadz M, Lu S, et al. The conserved domain database in 2023. Nucleic Acids Research. 2023;51(D1): D384–D388. 10.1093/nar/gkac1096.

42. Cury J, Abby SS, Doppelt-Azeroual O, Néron B, Rocha EPC. Identifying Conjugative Plasmids and Integrative Conjugative Elements with CONJscan. In: De La Cruz F (ed.) Horizontal Gene Transfer. New York, NY: Springer US; 2020. p. 265–283. 10.1007/978-1-4939-9877-7_19.

43. Néron B, Denise R, Coluzzi C, Touchon M, Rocha EPC, Abby SS. MacSyFinder v2: Improved modelling and search engine to identify molecular systems in genomes. Peer Community Journal. 2023;3: e28. 10.24072/pcjournal.250.

44. Taboada B, Estrada K, Ciria R, Merino E. Operon-mapper: a web server for precise operon identification in bacterial and archaeal genomes. Hancock J (ed.) Bioinformatics. 2018;34(23): 4118–4120. 10.1093/bioinformatics/bty496.

45. Guglielmini J, Néron B, Abby SS, Garcillán-Barcia MP, La Cruz FD, Rocha EPC. Key components of the eight classes of type IV secretion systems involved in bacterial conjugation or protein secretion. Nucleic Acids Research. 2014;42(9): 5715–5727. 10.1093/nar/gku194.

46. Peeters N, Carrère S, Anisimova M, Plener L, Cazalé AC, Genin S. Repertoire, unified nomenclature and evolution of the Type III effector gene set in the *Ralstonia solanacearum* species complex. BMC Genomics. 2013;14(1): 859. 10.1186/1471-2164-14-859.

47. Aoun N, Georgoulis SJ, Avalos JK, Grulla KJ, Miqueo K, Tom C, et al. A pangenomic atlas reveals eco-evolutionary dynamics that shape type VI secretion systems in plant-pathogenic *Ralstonia*. Ruby EG, Allen C (eds) mBio. 2024;15(10): e00323-24. 10.1128/mbio.00323-24.

48. Shaffer M, Borton MA, Bolduc B, Faria JP, Flynn RM, Ghadermazi P, et al. kb_DRAM: annotation and metabolic profiling of genomes with DRAM in KBase. Marschall T (ed.) Bioinformatics. 2023;39(4): btad110. 10.1093/bioinformatics/btad110.

49. Zhang H, Yohe T, Huang L, Entwistle S, Wu P, Yang Z, et al. dbCAN2: a meta server for automated carbohydrate-active enzyme annotation. Nucleic Acids Research. 2018;46(W1): W95–W101. 10.1093/nar/gky418.

50. Faria JP, Liu F, Edirisinghe JN, Gupta N, Seaver SMD, Freiburger AP, et al. ModelSEED v2: High-throughput genome-scale metabolic model reconstruction with enhanced energy biosynthesis pathway prediction. bioRxiv. 2023; 10.1101/2023.10.04.556561.

51. Dziurzynski M, Gorecki A, Decewicz P, Ciuchcinski K, Dabrowska M, Dziewit L. Development of the LCPDb-MET database facilitating selection of PCR primers for the detection of metal metabolism and resistance genes in bacteria. Ecological Indicators. 2022;145: 109606. 10.1016/j.ecolind.2022.109606.

52. Garcillán-Barcia MP, Redondo-Salvo S, Vielva L, De La Cruz F. MOBscan: Automated Annotation of MOB Relaxases. In: De La Cruz F (ed.) Horizontal Gene Transfer. New York, NY: Springer US; 2020. p. 295–308. 10.1007/978-1-4939-9877-7_21.

53. Pritchard L, Glover RH, Humphris S, Elphinstone JG, Toth IK. Genomics and taxonomy in diagnostics for food security: soft-rotting enterobacterial plant pathogens. Analytical Methods. 2016;8: 12–24. 10.1039/C5AY02550H.

54. Morpheus. https://software.broadinstitute.org/morpheus

55. Weisberg AJ, Chang JH. Mobile Genetic Element Flexibility as an Underlying Principle to Bacterial Evolution. Annual Review of Microbiology. 2023;77(1): 603–624. 10.1146/annurev-micro-032521-022006.

56. Hullahalli K, Rodrigues M, Nguyen UT, Palmer K. An Attenuated CRISPR-Cas System in *Enterococcus faecalis* Permits DNA Acquisition. Kline KA (ed.) mBio. 2018;9(3): e00414–18. 10.1128/mBio.00414-18.

57. Gonçalves OS, De Queiroz MV, Santana MF. Potential evolutionary impact of integrative and conjugative elements (ICEs) and genomic islands in the *Ralstonia solanacearum* species complex. Scientific Reports. 2020;10(1): 12498. 10.1038/s41598-020-69490-1.

58. Gonçalves OS, Campos KF, De Assis JCS, Fernandes AS, Souza TS, Do Carmo Rodrigues LG, et al. Transposable elements contribute to the genome plasticity of Ralstonia solanacearum species complex. Microbial Genomics. 2020;6(5). 10.1099/mgen.0.000374.

59. Gonçalves OS, Souza FDO, Bruckner FP, Santana MF, Alfenas-Zerbini P. Widespread distribution of prophages signaling the potential for adaptability and pathogenicity evolution of *Ralstonia solanacearum* species complex. Genomics. 2021;113(3): 992–1000. 10.1016/j.ygeno.2021.02.011.

60. Guidot A, Coupat B, Fall S, Prior P, Bertolla F. Horizontal gene transfer between *Ralstonia solanacearum* strains detected by comparative genomic hybridization on microarrays. The ISME Journal. 2009;3(5): 549–562. 10.1038/ismej.2009.14.

61. Wicker E, Lefeuvre P, De Cambiaire JC, Lemaire C, Poussier S, Prior P. Contrasting recombination patterns and demographic histories of the plant pathogen *Ralstonia solanacearum* inferred from MLSA. The ISME Journal. 2012;6(5): 961–974. 10.1038/ismej.2011.160.

62. Castillo JA. Analysis of the speciation process suggests a dual lifestyle in the plant pathogen *Ralstonia solanacearum* species complex. European Journal of Plant Pathology. 2023;166(2): 251–257. 10.1007/s10658-023-02670-7.

63. Li P, Wang D, Yan J, Zhou J, Deng Y, Jiang Z, et al. Genomic Analysis of Phylotype I Strain EP1 Reveals Substantial Divergence from Other Strains in the *Ralstonia solanacearum* Species Complex. Frontiers in Microbiology. 2016;7: 1719. 10.3389/fmicb.2016.01719.

64. Li X, Huang X, Chen G, Zou L, Wei L, Hua J. Complete genome sequence of the sesame pathogen *Ralstonia solanacearum* strain SEPPX 05. Genes & Genomics. 2018;40(6): 657–668. 10.1007/s13258-018-0667-3.

65. Das A, Xie YH. The *Agrobacterium* T-DNA Transport Pore Proteins VirB8, VirB9, and VirB10 Interact with One Another. Journal of Bacteriology. 2000;182(3): 758–763. 10.1128/JB.182.3.758-763.2000.

66. Christie PJ. Type IV secretion: the *Agrobacterium* VirB/D4 and related conjugation systems. Biochimica et Biophysica Acta. 2004;1694(1–3): 219–234. 10.1016/j.bbamcr.2004.02.013.

67. Groenewold MK, Hebecker S, Fritz C, Czolkoss S, Wiesselmann M, Heinz DW, et al. Virulence of *Agrobacterium tumefaciens* requires lipid homeostasis mediated by the lysyl-phosphatidylglycerol hydrolase AcvB. Molecular Microbiology. 2019;111(1): 269–286. 10.1111/mmi.14154.

68. González ET, Brown DG, Swanson JK, Allen C. Using the *Ralstonia solanacearum* Tat Secretome To Identify Bacterial Wilt Virulence Factors. Applied and Environmental Microbiology. 2007;73(12): 3779–3786. 10.1128/AEM.02999-06.

69. Dugelay C, Ferrarin S, Terradot L. Crystal structure of the virulence protein J (VirJ) domain 1 from *Brucella abortus*. *Acta Crystallographica Section F*, Structural Biology Communications. 2025;F81(Pt 9): 374–380. 10.1107/S2053230X25006697.

70. Pantoja M, Chen L, Chen Y, Nester EW. *Agrobacterium* type IV secretion is a two-step process in which export substrates associate with the virulence protein VirJ in the periplasm. Molecular Microbiology. 2002;45(5): 1325–1335. 10.1046/j.1365-2958.2002.03098.x.

71. Austin S, Nordström K. Partition-mediated incompatibility of bacterial plasmids. Cell. 1990;60(3): 351–354. 10.1016/0092-8674(90)90584-2.

72. Ravenhall M, Škunca N, Lassalle F, Dessimoz C. Inferring Horizontal Gene Transfer. Wodak S (ed.) PLOS Computational Biology. 2015;11(5): e1004095. 10.1371/journal.pcbi.1004095.

73. Macho AP. Subversion of plant cellular functions by bacterial type-III effectors: beyond suppression of immunity. New Phytologist. 2016;210(1): 51–57. 10.1111/nph.13605.

74. Tan X, Qiu H, Li F, Cheng D, Zheng X, Wang B, et al. Complete Genome Sequence of Sequevar 14M *Ralstonia solanacearum* Strain HA4-1 Reveals Novel Type III Effectors Acquired Through Horizontal Gene Transfer. Frontiers in Microbiology. 2019;10: 1893. 10.3389/fmicb.2019.01893.

75. Genin S, Boucher C. Lessons Learned from the Genome Analysis of *Ralstonia solanacearum*. Annual Review of Phytopathology. 2004;42(1): 107–134. 10.1146/annurev.phyto.42.011204.104301.

76. Evseeva D, Pecrix Y, Kucka M, Weiler C, Franzl C, Vlková-Žlebková M, et al. Emergence and host range expansion of an epidemic lineage of *Ralstonia solanacearum*. bioRxiv. 2024; 10.1101/2024.10.16.618685.

77. Prokchorchik M, Pandey A, Moon H, Kim W, Jeon H, Jung G, et al. Host adaptation and microbial competition drive *Ralstonia solanacearum* phylotype I evolution in the Republic of Korea. Microbial Genomics. 2020;6(11). 10.1099/mgen.0.000461.

78. Nash ZM, Cotter PA. *Bordetella* Filamentous Hemagglutinin, a Model for the Two-Partner Secretion Pathway. Sandkvist M, Cascales E, Christie PJ (eds) Microbiology Spectrum. 2019;7(2): PSIB-0024-2018. 10.1128/microbiolspec.psib-0024-2018.

79. Hodak H, Clantin B, Willery E, Villeret V, Locht C, Jacob-Dubuisson F. Secretion signal of the filamentous haemagglutinin, a model two-partner secretion substrate. Molecular Microbiology. 2006;61(2): 368–382. 10.1111/j.1365-2958.2006.05242.x.

80. Pérez A, Merino M, Rumbo-Feal S, Álvarez-Fraga L, Vallejo JA, Beceiro A, et al. The FhaB/FhaC two-partner secretion system is involved in adhesion of *Acinetobacter baumannii* AbH12O-A2 strain. Virulence. 2017;8(6): 959–974. 10.1080/21505594.2016.1262313.

81. Trouillon J, Attrée I, Elsen S. The regulation of bacterial two-partner secretion systems. Molecular Microbiology. 2023;120(2): 159–177. 10.1111/mmi.15112.

82. Jacob-Dubuisson F, Guérin J, Baelen S, Clantin B. Two-partner secretion: as simple as it sounds? Research in Microbiology. 2013;164(6): 583–595. 10.1016/j.resmic.2013.03.009.

83. Guérin J, Bigot S, Schneider R, Buchanan SK, Jacob-Dubuisson F. Two-Partner Secretion: Combining Efficiency and Simplicity in the Secretion of Large Proteins for Bacteria-Host and Bacteria-Bacteria Interactions. Frontiers in Cellular and Infection Microbiology. 2017;7: 148. 10.3389/fcimb.2017.00148.

84. Leo JC, Grin I, Linke D. Type V secretion: mechanism(s) of autotransport through the bacterial outer membrane. Philosophical Transactions of the Royal Society B: Biological Sciences. 2012;367: 1088–1101. 10.1098/rstb.2011.0208.

85. Mathivanan K, Chandirika JU, Vinothkanna A, Yin H, Liu X, Meng D. Bacterial adaptive strategies to cope with metal toxicity in the contaminated environment – A review. Ecotoxicology and Environmental Safety. 2021;226: 112863. 10.1016/j.ecoenv.2021.112863.

86. Pal A, Bhattacharjee S, Saha J, Sarkar M, Mandal P. Bacterial survival strategies and responses under heavy metal stress: a comprehensive overview. Critical Reviews in Microbiology. 2022;48(3): 327–355. 10.1080/1040841X.2021.1970512.

87. Coupat B, Chaumeille-Dole F, Fall S, Prior P, Simonet P, Nesme X, et al. Natural transformation in the *Ralstonia solanacearum* species complex: number and size of DNA that can be transferred. FEMS Microbiology Ecology. 2008;66(1): 14–24. 10.1111/j.1574-6941.2008.00552.x.

88. Weisberg AJ, Wu Y, Chang JH, Lai EM, Kuo CH. Virulence and Ecology of Agrobacteria in the Context of Evolutionary Genomics. Annual Review of Phytopathology. 2023;61(1): 1–23. 10.1146/annurev-phyto-021622-125009.

89. Shin GY, Asselin JA, Smith A, Aegerter B, Coutinho T, Zhao M, et al. Plasmids encode and can mobilize onion pathogenicity in *Pantoea agglomerans*. The ISME Journal. 2025;19(1): wraf019. 10.1093/ismejo/wraf019.

90. Banerjee G, Chatterjee S, Banerjee P, Chattopadhyay P. Diversity of Secretion System Apparatus in Tomato Wilt Causing *Ralstonia solanacearum* Strains: a Comparative Analysis Using *in-silico* Approach. bioRxiv. 2022; 10.1101/2022.04.21.489029.

91. Ding S, Yu L, Lan G, Tang Y, Li Z, He Z, et al. Identification and genomic characterization of *Ralstonia pseudosolanacearum* strains isolated from pepino melon in China. Physiological and Molecular Plant Pathology. 2023;125: 101977. 10.1016/j.pmpp.2023.101977.

92. Ding S, Ma Z, Yu L, Lan G, Tang Y, Li Z, et al. Comparative genomics and host range analysis of four *Ralstonia pseudosolanacearum* strains isolated from sunflower reveals genomic and phenotypic differences. BMC Genomics. 2024;25(1): 191. 10.1186/s12864-024-10087-7.

93. Salanoubat M, Genin S, Artiguenave F, Gouzy J, Mangenot S, Arlat M, et al. Genome sequence of the plant pathogen *Ralstonia solanacearum*. Nature. 2002;415: 497–502. 10.1038/415497a.

94. Remenant B, Coupat-Goutaland B, Guidot A, Cellier G, Wicker E, Allen C, et al. Genomes of three tomato pathogens within the *Ralstonia solanacearum* species complex reveal significant evolutionary divergence. BMC Genomics. 2010;11: 379. 10.1186/1471-2164-11-379.

95. Morales VM, Sequeira L. Indigenous Plasmids in *Pseudomonas solanacearum*. Phytopathology. 1985;75(7): 767–771. 10.1094/Phyto-75-767.

96. Hamilton HL, Domínguez NM, Schwartz KJ, Hackett KT, Dillard JP. *Neisseria gonorrhoeae* secretes chromosomal DNA via a novel type IV secretion system. Molecular Microbiology. 2005;55(6): 1704–1721. 10.1111/j.1365-2958.2005.04521.x.

97. Callaghan MM, Heilers JH, Van Der Does C, Dillard JP. Secretion of Chromosomal DNA by the *Neisseria gonorrhoeae* Type IV Secretion System. In: Backert S, Grohmann E (eds) Type IV Secretion in Gram-Negative and Gram-Positive Bacteria. Springer International Publishing; 2017. p. 323–345. 10.1007/978-3-319-75241-9_13.

98. Jurcisek JA, Brockman KL, Novotny LA, Goodman SD, Bakaletz LO. Nontypeable *Haemophilus influenzae* releases DNA and DNABII proteins via a T4SS-like complex and ComE of the type IV pilus machinery. Proceedings of the National Academy of Sciences. 2017;114(32): E6632–E6641. 10.1073/pnas.1705508114.

99. Goodman SD, Scocca JJ. Identification and arrangement of the DNA sequence recognized in specific transformation of *Neisseria gonorrhoeae*. Proceedings of the National Academy of Sciences. 1988;85(18): 6982–6986. 10.1073/pnas.85.18.6982.

100. Dewar AE, Hao C, Belcher LJ, Ghoul M, West SA. Bacterial lifestyle shapes pangenomes. Proceedings of the National Academy of Sciences. 2024;121(21): e2320170121. 10.1073/pnas.2320170121.

