## Supplemental Figures for "Lifestyle-associated variation in type IV secretion systems between phytopathogenic and environmental *Ralstonia*"

In addition to the five supplemental figures, a Zenodo repository contains supplemental tables and files like multiple sequence alignments, phylogenetic trees, and GenBank files that can be used for clinker synteny analyses (DOI: 10.5281/zenodo.14873518).

---

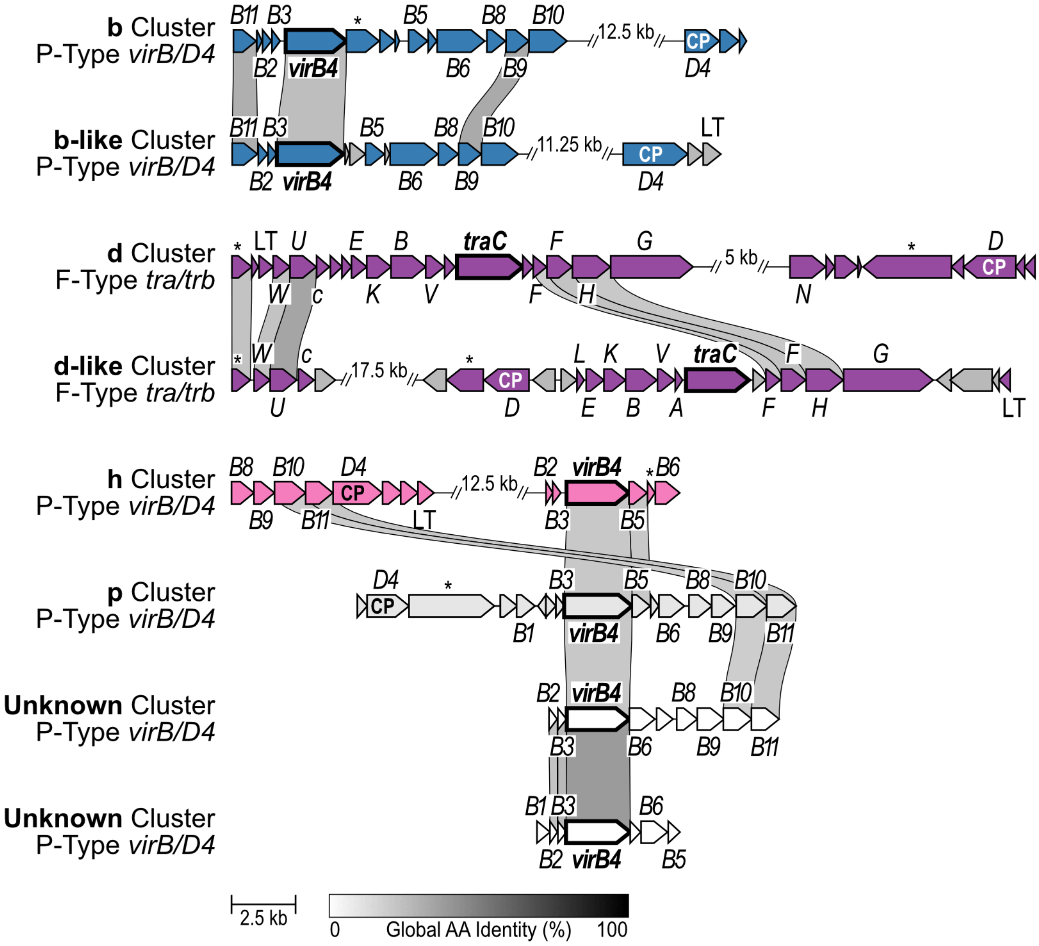


**Figure S1.** Synteny comparison of T4SS gene clusters: b vs. b-like, d vs. d-like, and h and p vs. the two unknown clusters. The designation of "b-like" and "d-like" was assigned to clusters that had conserved, but low identity, gene order with the reference cluster. The linkages between gene arrows were drawn with clinker software, which performs pairwise all-vs-all global amino acid identity comparisons with a threshold of 30% amino acid identity. Genes in each cluster are labeled based on F/P/I/GI-type classification conventions: *tra*/*trb* (F-type) and *virB*/*D4* (P-type). The *virB4* homologs have a bolded name (*traC* and *virB4*) and a bolded gene arrow border. The genes annotated as the type IV coupling protein are labeled "CP". "LT" indicates genes annotated as lytic transglycosylase genes. Genes marked with an asterisk (*) are T4SS genes that were not able to be annotated within the indicated F/P/I/GI-type convention. Gene clusters were defined using NCBI RefSeq annotations and NCBI Conserved Domain Database. Gaps are displayed where the cluster had regions without predicted T4SS genes; patterns of non-T4SS genes are not always consistent among all members of each cluster. Visualization adjustments were made using Affinity Designer.

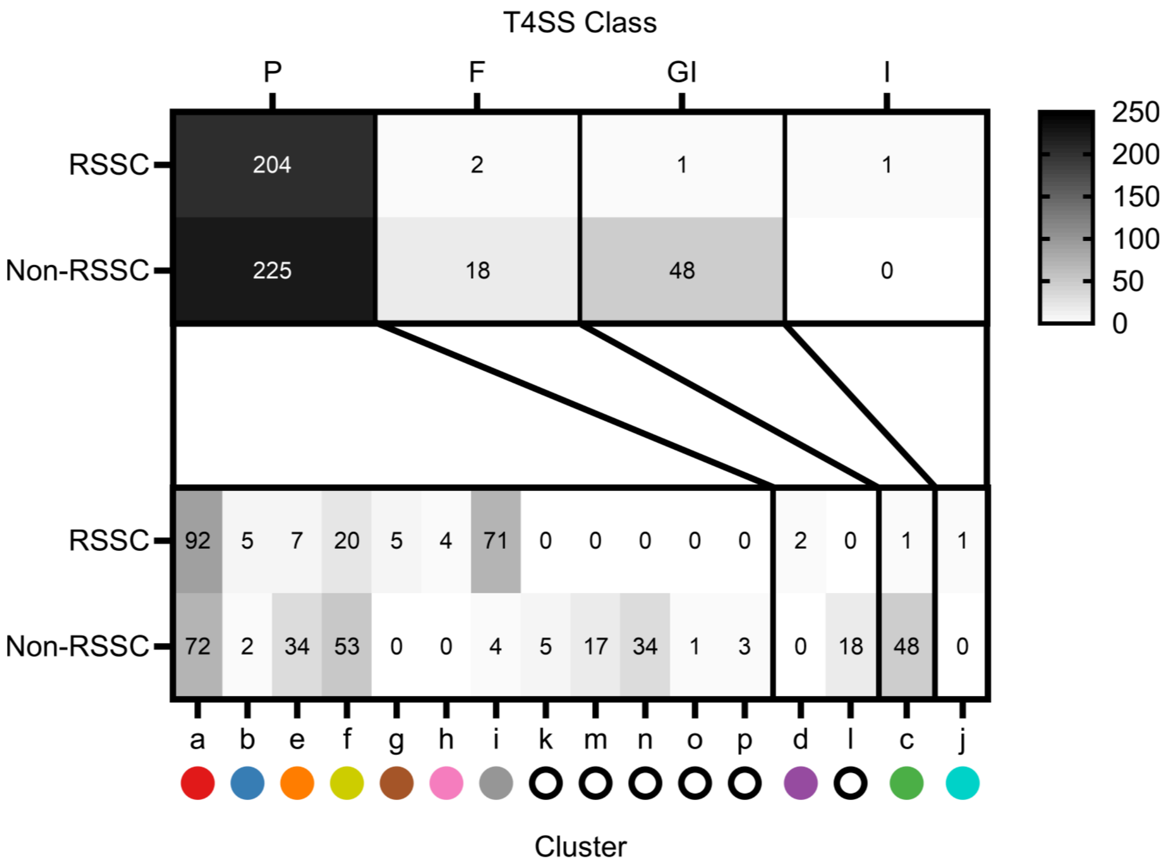


**Figure S2.** Distribution of T4SS gene clusters by T4SS class using F/P/I/GI-type classification nomenclature. The majority of the T4SS gene clusters found in *Ralstonia* genomes are P-type. Numbers indicate the number of T4SS gene clusters of that cluster/class in the RSSC pathogens vs. non-RSSC environments. The counts of T4SS gene clusters are based on the number of *virB4* genes in each category, including 494 of the 496 identified by the HMMER and BLASTp searches, as well as the five manually identified sequences. The two T4SS clusters excluded are the two unknown clusters.

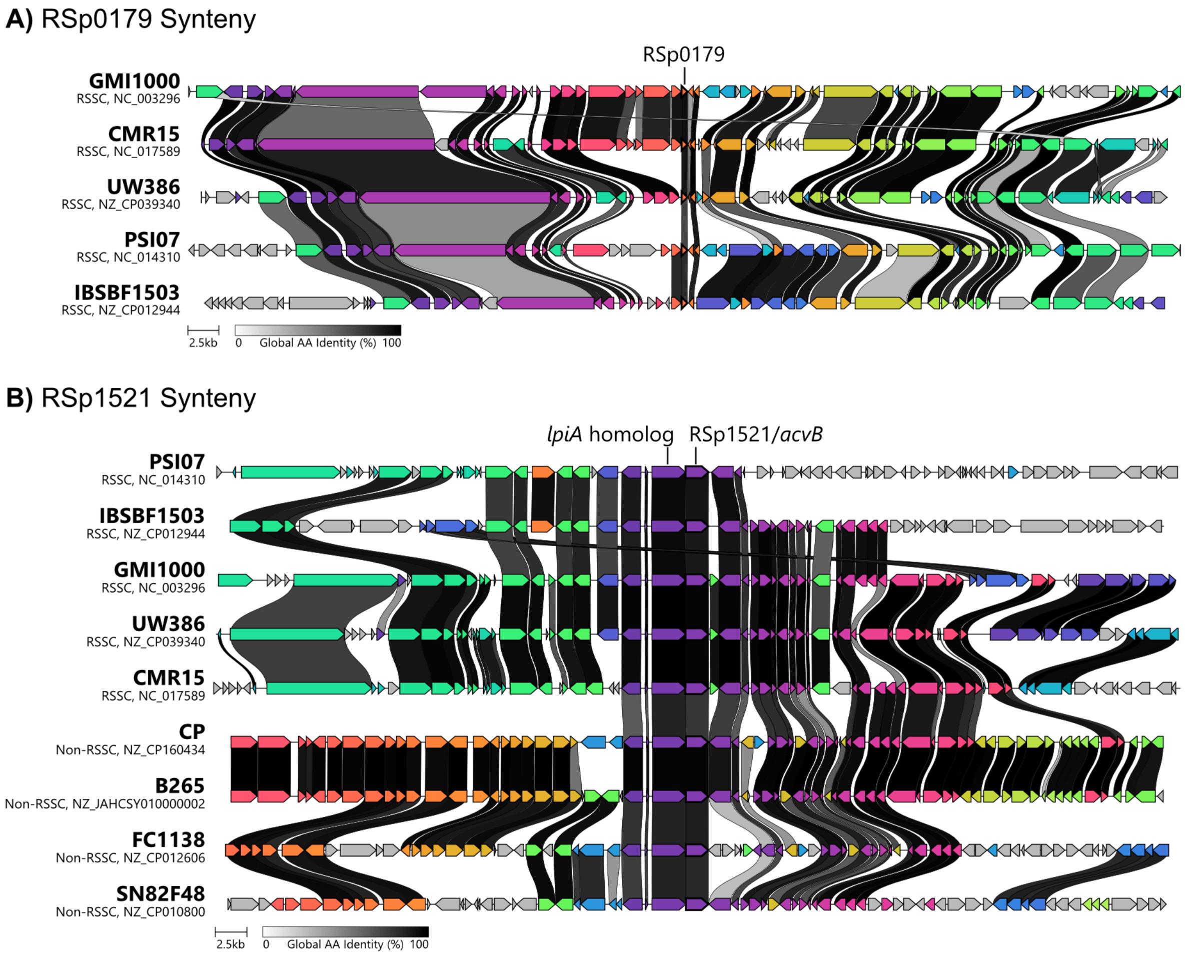


**Figure S3.** Synteny analysis of the gene neighborhoods of two genes that are unlikely to be T4SS-associated: **A)** RSp0179 and **B)** RSp1521. Both genes were consistently found on the megaplasmid. Gene arrow colors are assigned by clinker and genes of the same exact color are homologous, other than gray genes, which do not have homologs in this depiction. The linkages between gene arrows are shaded based on global amino acid identity. The synteny analyses were performed using clinker. Visualization adjustments were made using Affinity Designer.

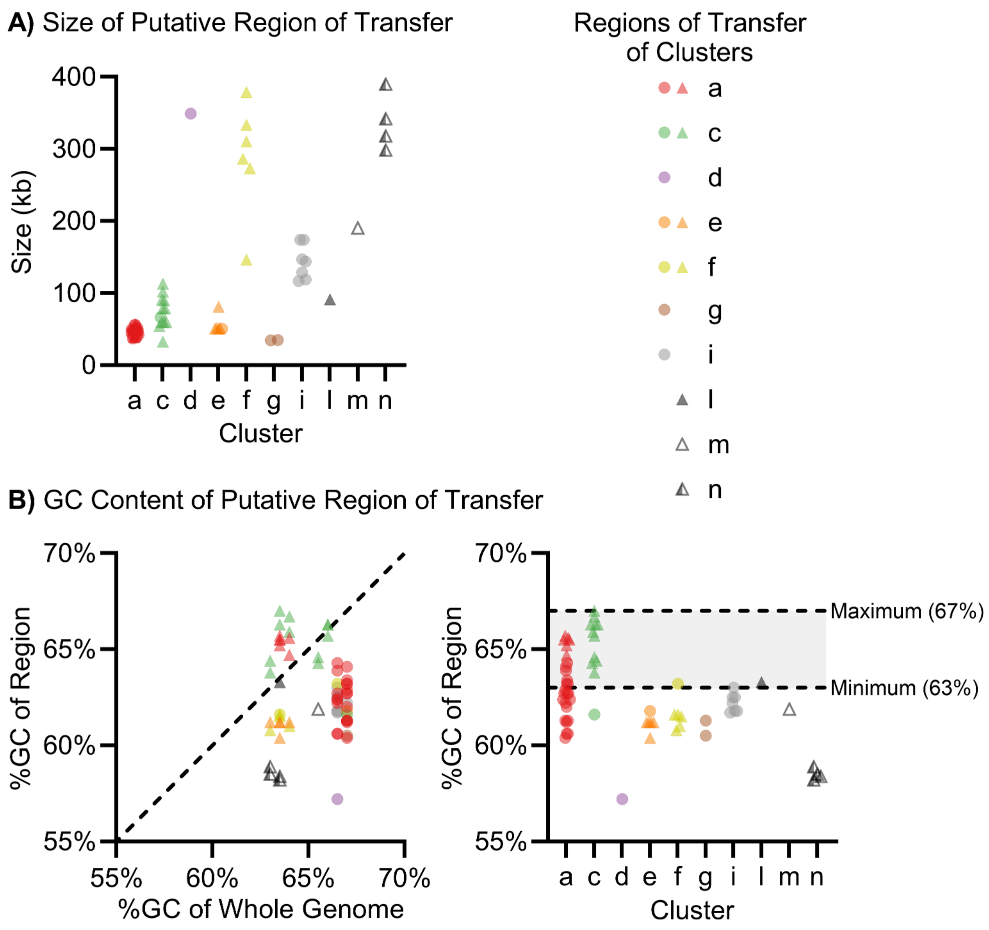


**Figure S4.** Properties of the regions of transfer. **A)** Sizes of putative regions of transfer. The size in kilobases is shown on the y-axis and the cluster is on the x-axis. Each point represents one region of transfer. Circles indicate RSSC phytopathogens and triangles indicate non-RSSC environmentals. **B)** GC content of putative regions of transfer. Left: the GC content of the whole genome on the x-axis and the GC content of only the putative region of transfer on the y-axis. The dashed line represents a one to one relationship between the GC content of the region and the GC content of the whole genome. Each point represents one region of transfer in one genome. Circles indicate RSSC phytopathogens and triangles indicate non-RSSC environmentals. Right: the GC content of the putative region of transfer on the y-axis and the cluster on the x-axis. The dashed lines labeled “Maximum (67%)” and “Minimum (63%)” represent the maximum and minimum GC content of the whole genomes used in this analysis. Each point represents one region of transfer. Circles indicate RSSC phytopathogens and triangles indicate non-RSSC environmentals.

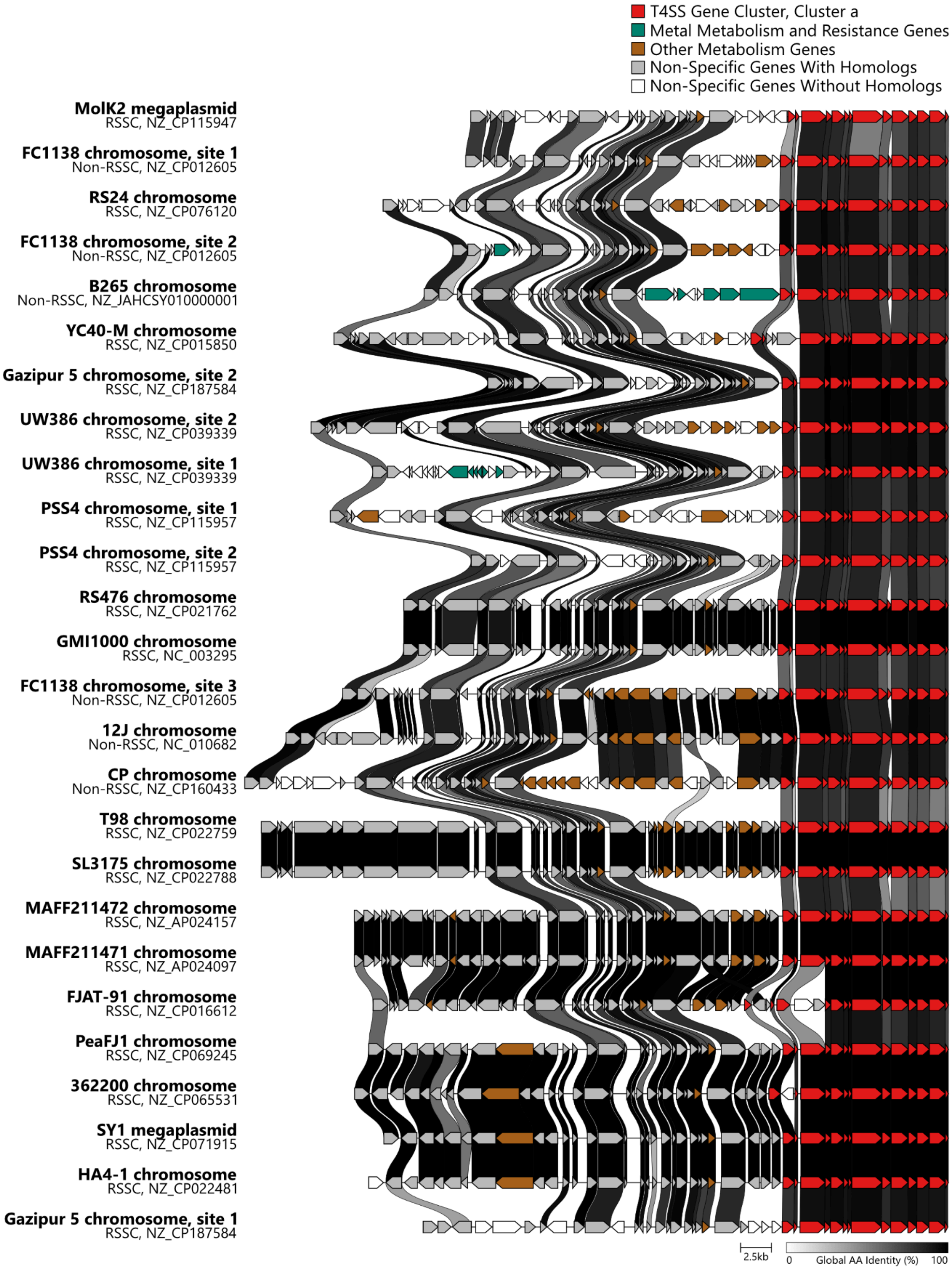


**Figure S5.** Example of cargo gene analysis using cluster a. The genes in T4SS cluster a are red, the heavy metal metabolism and resistance genes are green, and the other metabolism genes are brown. Genes that do not fall into the selected cargo categories are gray or white. Gray genes are those that have homologs among the genes analyzed. White genes are those that do not have homologs present in this analysis. The synteny analysis was performed using clinker. Gene colors were edited using Affinity Designer. Interactive HTML files for other putative region of transfer visualizations like this for the other clusters with complete genomes are available in Supplemental Files S27-S36.
